# The developmental cycle of *Dictyostelium discoideum* ensures curing of a mycobacterial infection at both cell-autonomous level and by collaborative exclusion

**DOI:** 10.1101/586263

**Authors:** Ana Teresa López-Jiménez, Monica Hagedorn, Matthieu J. Delincé, John McKinney, Thierry Soldati

## Abstract

During its life cycle, the social amoeba *Dictyostelium discoideum* alternates between a predatory amoeba and a facultative multicellular form. The single-celled amoeba is a well-established model system to study cell-autonomous mechanisms of phagocytosis and defence against intracellular bacterial pathogens, whereas the multicellular forms are arising as models to study the emergence of innate immune defence strategies. Importantly, during evolution, prokaryotes have also evolved their own strategies to resist predation. Considering these complex ecological relationships, we wondered whether *D. discoideum* cells infected with intracellular pathogenic mycobacteria would be able to undergo their developmental cycle and what would be the fate of the infection. We show that the combination of cell-autonomous mechanisms and the organisation into a multicellular organism leads to the efficient multistep-curing of a mycobacteria-infected population, thereby ensuring germ-free spores and progeny. Specifically, using a microfluidic device to trap single infected cells, we revealed that in the first curing phase, individual cells rely on three mechanisms to release intracellular bacteria: exocytic release, ejection and lytic release. The second phase occurs at the collective level, when remaining infected cells are excluded from the forming cell aggregates.

## INTRODUCTION

During the various phases of its life cycle, the social amoeba *Dictyostelium discoideum* alternates between a single-celled phagocytic amoeba and a higher order multicellular form. Such facultative multicellular organisms have the intriguing ability to adjust their life-form to environmental conditions. Indeed, in the soil, the single-celled amoebae feed on bacteria by phagocytosis. It encounters and predates on a wide range of bacteria species but it is also exposed to periods of scarcity. During which hundreds of thousands of amoebae undergo a complex developmental program that gives rise to a multicellular organism (Loomis, 2015; Raper, 1935; Weijer, 2004).

Because of this duality, *D. discoideum* has emerged as a powerful model to study both cell-autonomous defence processes as well as the mechanisms that ensure innate immunity at the multicellular stage.

### Cell autonomous defence against bacteria

In vegetative growing cells, *D. discoideum* uses similar bactericidal tools as phagocytes of the innate immune system of animals to fight infection (Boulais et al., 2010). This includes acidification (Marchetti et al., 2009), lysosomal hydrolases (Journet et al., 1999), NOX-ROS (Lardy et al., 2005; Zhang and Soldati, 2013), a role for Mg^2+^ (Lelong et al., 2011) and Fe^2+^ (Bozzaro et al., 2013). A complete autophagy pathway also contributes to the control of some pathogens that damage their vacuole and/or escape to the cytosol (Cardenal-Munoz et al., 2017; Huang and Brumell, 2009). As shown previously, *D. discoideum* can be efficiently infected by the fish and frog pathogen *M. marinum*, which is genetically closely related to *M. tuberculosis* (Stinear et al., 2008). Initially, after phagocytosis by *D. discoideum, M. marinum* hampers the acidification and maturation of the compartment where it resides, and establishes a permissive niche where the bacteria can survive and replicate (Hagedorn and Soldati, 2007; Solomon et al., 2003). At later stages of infection, the pathogen translocates into the host cell cytosol (Cardenal-Munoz et al., 2017; Hagedorn and Soldati, 2007), a process dependent on the mycobacterial secretion locus ESX-1 (Cardenal-Munoz et al., 2017). During escape to the cytosol, autophagy appears to play a major and complex role, and autophagy null mutants are incapacitated in controlling the *M. marinum* infection (Cardenal-Munoz et al., 2017). Finally, the cytosolic bacterium is released from their host by cell lysis, or it is non-lytically ejected through an actin-dependent egress-structure, the ejectosome (Gerstenmaier et al., 2015; Hagedorn et al., 2009). Most of these stages have been studied at the bulk population level, but it has recently emerged that host-pathogen interactions are sometimes dominated by stochastic events that rely on the phenotypic heterogeneity within a population (Dhar et al., 2016). Hence, quantitative single-cell approaches start to be prevalent to study the biological processes involved during infection. These methodologies include the utilization of microfluidic devices to encapsulate single infected cells, which can be visualized for long periods of time. In that regard, a new microfluidic device, the InfectChip, was specifically designed to monitor amoeba-bacteria interactions and was instrumental to reveal and understand the complexity of the infection process (Delince et al., 2016).

### Emergence of innate immunity

When local sources of food are consumed, hundreds of thousands of amoebae undergo a complex developmental program that gives rise to a multicellular organism (Loomis, 2015; Raper, 1935; Weijer, 2004). Interestingly, this form of “facultative” multicellularity coincides with the invention of self-nonself recognition via allelic pairs of polymorphic receptors (Benabentos et al., 2009; Hirose et al., 2011; Strassmann, 2016). The process starts with the establishment of concentric waves of extracellular cAMP that serves as a chemoattractant for individual cells (Konijn et al., 1967; Loomis, 2008). Simultanously, *D. discoideum* begins to express adhesion molecules at the cell surface (such as Contact Site A, CsA) (Gerisch, 1968; Ochiail et al., 1982) to gather together in compact streams while migrating to the center of the cAMP gradient, where they form an early aggregate. This aggregate gives rise to the slug, a true thermo- and phototactic multicellular organism (Raper, 1935). The slug is motile (Poff et al., 1973), externally delimited by a cellulose sheath (Freeze and Loomis, 1977) and it contains differentiating and specialized cells (Wang et al., 1988), which implies cell differentiation and division of tasks. At this stage, cells are already committed to form the stalk or the spore mass in the future fruiting body. In addition, the Sentinel cells (S-cells) are the only specialized cells which retain the ability to phagocytose, and patrol the slug in search of particles or bacteria to be eliminated (Chen et al., 2007). This process is proposed as the evolutionary origin of the metazoan immune system. Furthermore, S-cells have been shown to extrude DNA-based extracellular traps (ET) to immobilize and kill bacteria, similarly to neutrophils and other granulocytes (Zhang et al., 2016). Finally, slugs transition into fruiting bodies (Loomis, 2015; Raper, 1935), composed of a sorus filled with spores mounted on a stalk of dead cells. Spores are much more resistant to environmental stresses than cells and can remain dormant for decades. When conditions ameliorate again, single-cell amoebae germinate from the spores and the cycle is closed.

While most wild *D. discoideum* isolates engage into their developmental program only when the bacteria supply is exhausted, others start aggregation when some bacteria remain. In addition, these isolates tend to carry edible bacteria species during their multicellular stage and were thus named “farmers” (Brock et al., 2013). These bacteria would serve as food source for the newly germinated amoeba, which would confer an ecological advantage in harsh environments. As a drawback, farmer *D. discoideum* strains have been shown to migrate less as slugs and suffer from a reduction in spore viability. Recently, stable association of *D. discoideum* and bacteria from the genus *Burkholderia* has been shown to be responsible for the induction of the farmer phenotype in the amoeba (DiSalvo et al., 2015). *Bulkholderia* species drive their own persistent carriage through all stages of *D. discoideum* developmental cycle as symbionts (Shu et al., 2018) and also allow the secondary co-colonization by other edible bacteria, proposed as commensalism or mutualism (DiSalvo et al., 2015). While the farmer phenotype can be induced in naïve *D. discoideum* by the mere coculture with *Bulkholderia*, wild-isolates of *D. discoideum* have been shown to develop permanent adaptations to tolerate this endosymbiont (Shu et al., 2018). Although the precise mechanisms that enable permissibility to *Bulkholderia* are not know, the farmer phenotype has been associated to an increased secretion of extracellular lectins by *D. discoideum*, that may protect the bacteria or induce their tolerance (Dinh et al., 2018; DiSalvo et al., 2015; Shu et al., 2018).

### Organising the resistance

During evolution, as protozoa evolved as powerful bacteria predators, equipped with a collection of mechanisms for efficient chase, killing and digestion (Cosson and Lima, 2014), prokaryotes were under extreme selective pressure to acquire the necessary adaptations to resist predation. As a consequence, they developed different strategies for survival, from merely avoiding or resisting grazing to pathogenic traits such as toxin secretion (Matz and Kjelleberg, 2005). For this reason, amoebae constitute the training ground for many pathogens, which became subsequently equipped to subvert the bactericidal apparatus of the phagocytic cells of the animal immune system (Molmeret et al., 2005). This is the case of the human parasites *Legionella pneumophila* (Escoll et al., 2013) and *Vibrio cholerae* (Van der Henst et al., 2016), for which amoebae act also as a currently important ecological reservoir (Baker and Brown, 1994). Vice versa, evidence is also cumulating that animal and human pathogens such as mycobacterium species (e.g. *Mycobacterium avium, Mycobacterium marinum*) can survive and even replicate in environmental protozoa (Cirillo et al., 1997; Lamrabet et al., 2012a; Lamrabet et al., 2012b; Mba Medie et al., 2011; Salah et al., 2009). As a result, several amoeba species and in particular the experimentally tractable *D. discoideum* have emerged as a powerful model system to study bacterial infection (Bozzaro and Eichinger, 2011; Cosson and Soldati, 2008; Tosetti et al., 2014).

Here, we demonstrate that *D. discoideum* cells heavily infected with pathogenic *M. marinum* are able to cure the intracellular infection during their developmental cycle. We show that this occurs in two temporarily and mechanistically distinct phases: via bacteria release at the single-cell level at the onset of starvation, and by collaborative exclusion of the remaining infected cells at a later aggregation phase.

## RESULTS

### *D. discoideum* cures a *M. marinum* infection during its developmental cycle

The infection cycle of *M. marinum* during the vegetative stage of *D. discoideum* is intensely studied. On the other hand, the impact of the multicellular developmental cycle on the fate of an infection is unknown. Therefore, we monitored the effect of starvation onset on an amoebae population homogeneously infected with *M. marinum*.

After confirming with flow cytometry and fluorescence microscopy that a large proportion of *D. discoideum* cells was infected and no extracellular bacteria remained (Fig S1), the infected cells were switched to starving conditions at two different stages of infection, the early vacuolar stage (6 hpi; Fig 1A-D, Movie S1) and a later mainly cytosolic stage (24 hpi; Fig 1E-H). Surprisingly, even though the vast majority of cells were initially infected (Fig 1A, A’, E, E’ and Fig S1), and independently of the infection stage (vacuolar or cytosolic), *D. discoideum* cells followed their differentiation program to yield apparently uninfected sori. At the onset of starvation, the fluorescent bacteria (Fig 1A, A’, E, E’) colocalized with the *D. discoideum* cells visible in the brightfield image. However, bacteria seemed to be kept out from the streams at later phases of aggregation (Fig 1B, B’, F, F’). Indeed, the bacteria are found to outline the compact streams (Fig 1F’) observed in the widefield image (Fig 1F). Subsequently, the early culminants (Fig 1C, C’, F, F’) and the slug (Fig 1D, D’, G, G’) seemed to be free of any accumulation of fluorescent bacteria. Instead, the multicellular structures of hundreds of thousands of cells formed a clear lense through which fluorescent bacteria that have been left behind by the *D. discoideum* cells are observed on the agar surface. At the end of differentiation, spores are tightly packed into a sorus standing on a stalk of dead vacuolated cells (Fig 1H, H’). Free, fluorescent bacteria are observed to remain on the outer surface of this sorus, but are absent from the cells that make up the spherical cellulose-encased spore mass (Fig 2).

**Fig. 1.**
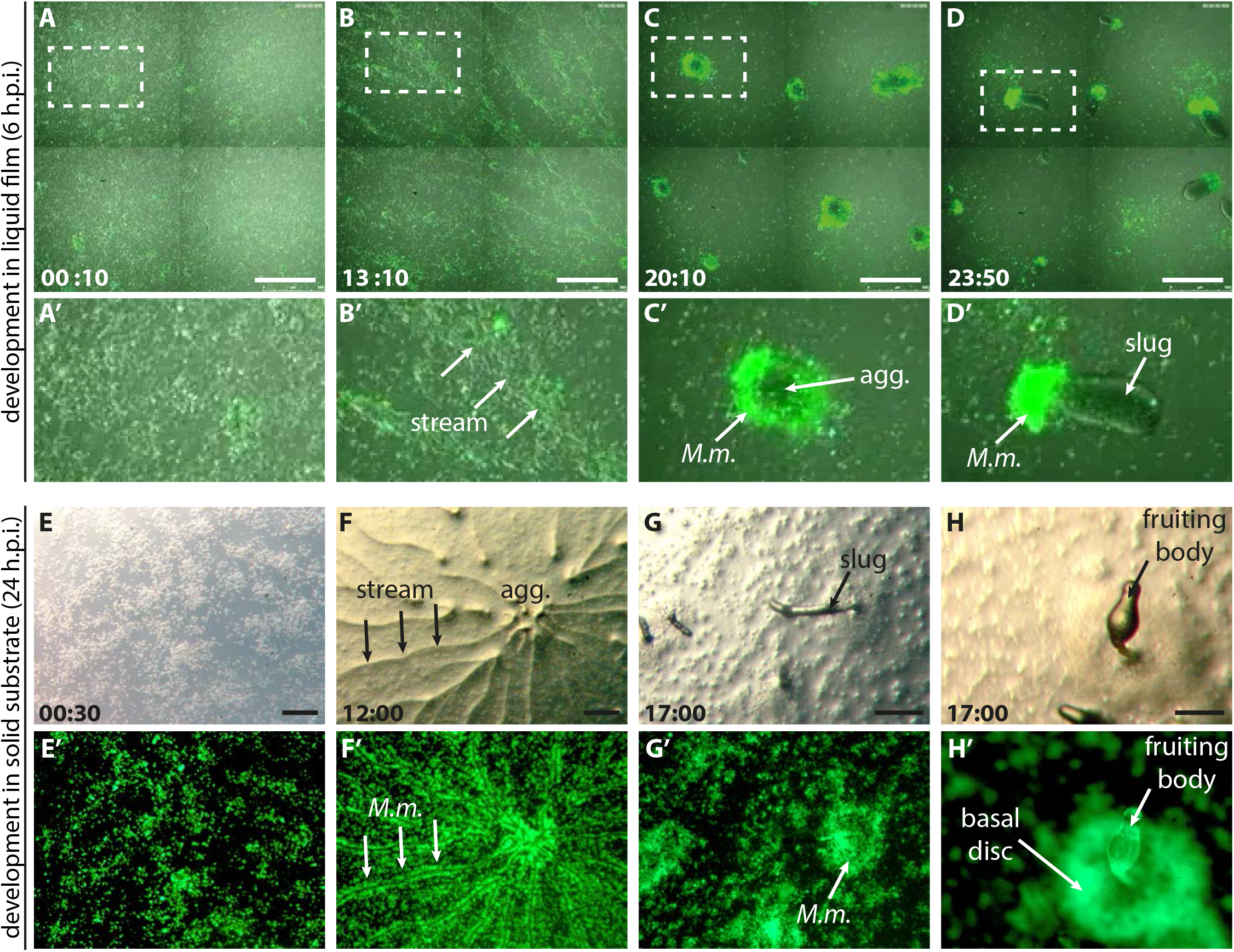
Upon differentiation, *D. discoideum* cells segregate from virulent *M. marinum*. **(A-D)** Infected *D. discoideum* cells were washed free of medium and covered with a thin film of non-nutrient buffer at 6 hpi, and imaged at 25° C every 10 minutes for 24 hours using a fluorescence widefield-microscope (*D. discoideum* visible in the brightfield images, *M. marinum* in green). 4 fields of view were stitched together (see also Movie S1). Time after starvation onset is indicated at the bottom left corner. Scale bars: 500 μm. (A’-D’) Areas at higher magnification are indicated with a dotted line. (E-H) Infected cells were washed free of medium after 24 hpi and transferred onto non-nutient-agar plates. The cells were left to develop at 25°C. Pictures were taken with a binocular microscope at the times indicated at the bottom left corner. The upper row presents the DIC images (E-H), the lower row shows the corresponding fluorescent images (E’-H’). Scale bars: 100 μm (E-F), 300 μm (G-H).

### The curing process is initiated at the single-cell stage during development

To follow the steps of the curing process in more detail, fluorescent *D. discoideum* cells expressing GFP-ABD were visualized during starvation. Macroscopically, the majority of the infected cells (Fig 2A) seemed to release their intracellular bacteria in the first 10 h after initiation of starvation, usually just before (Fig 2B) and during stream formation (Fig 2C see also Movie S1). While the cells moved towards the center of aggregation, fluorescent bacteria did not participate in the compact streams (Fig 2C).

**Fig. 2.**
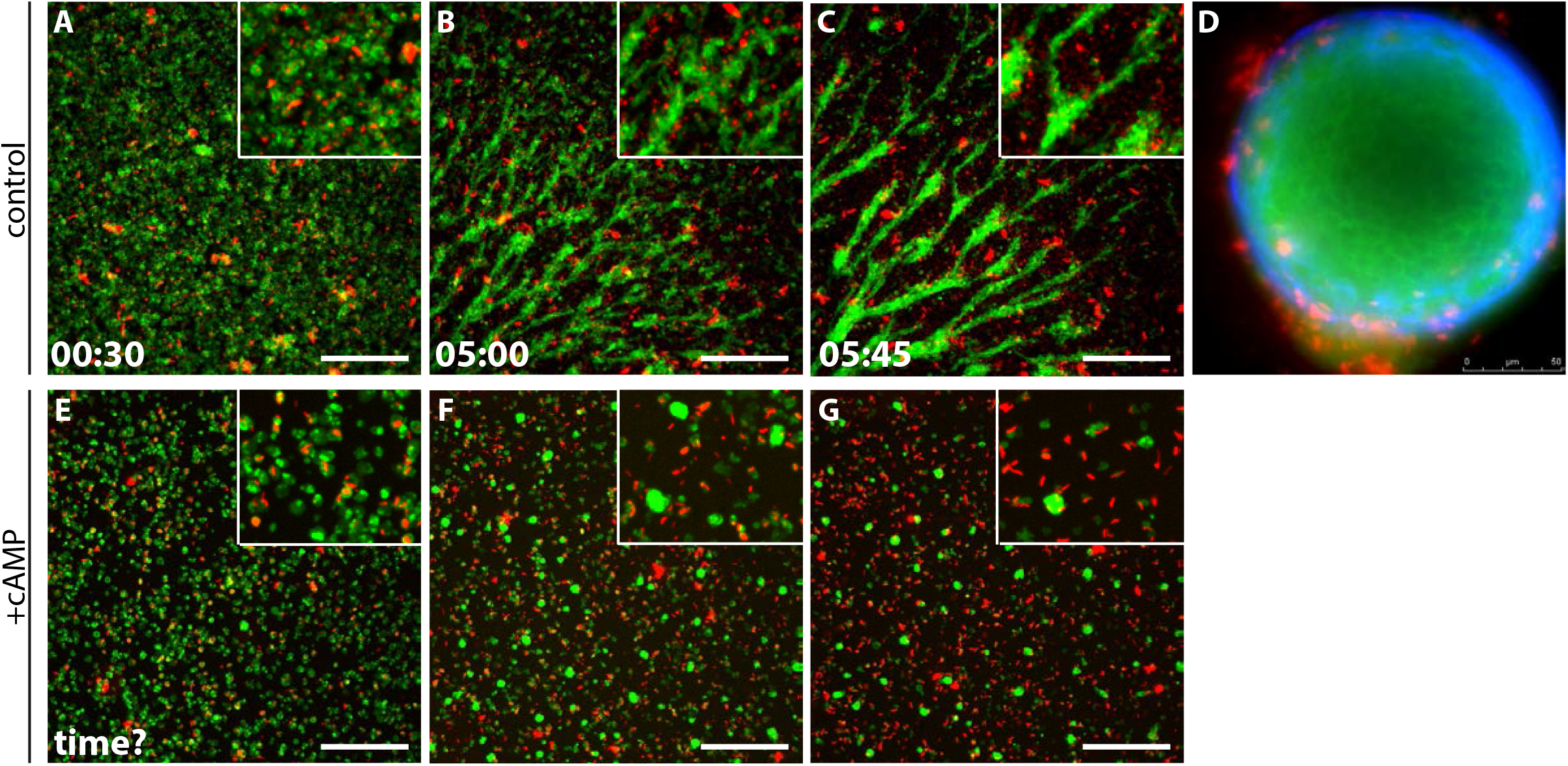
Curing of *M. marinum* infection is initiated at the single-cell level upon starvation. *D. discoideum* cells expressing GFP-ABD (green) were infected with mCherry *M. marinum* (red), and, at 6 hpi, were starved, in the absence (A-D) or presence (E-G) of exogenous cAMP (0.1 mM) and monitored by live-cell fluorescence microscopy. In the absence of exogenous cAMP, infected *D. discoideum* cells (A) became elongated and organized into streams moving towards the center of aggregation (B-C). Concomitantly with the start of streaming, a segregation was observed between *M. marinum* and *D. discoideum* cells. This separation becames more evident with time and increased with stream thickness and compaction (B–C, insets Movie S2, 12 frames/h). The final tight aggregate (D), ensheathed in cellulose (blue, stained with calcofluor) shows uninfected cells within the aggregate (green, stained with FM-4-64) and free bacteria (red) left out of the aggregate, outside the cellulose sheath. (E-G) Infected *D. discoideum* cells were starved at 6 hpi in the presence of excess exogenous cAMP (0.1 mM). While segregation between bacteria and host cells appeared to occur more slowly and not quite as efficiently after 5h (G), the formation of small aggregates of uninfected cells was nevertheless observed and a large number of extracellular bacteria were visible (G, inset). Scale bars: 100 μm.

In order to trigger tight multicellular streams and subsequently aggregate formation, *D. discoideum* cells secrete, relay and follow a gradient of cAMP and express developmentally-regulated cell-adhesion molecules. To decipher whether this early curing event is performed at the single-cell level, prior to aggregate formation, cells were starved in the presence of high concentrations of cAMP (Fig 2E-G). Under these conditions, multicellular streams and the formation of large aggregates are almost completely suppressed, but cells still express cell-adhesion molecules and form small aggregates by random collision. Live microscopy showed that initially infected *D. discoideum* cells assembled into small-sized aggregates. Interestingly, bacteria nevertheless still segregated from these structures (Fig 2F, G). This means that, even though streams are not formed, cells correctly initiated their developmental program and retained their capacity to keep out *M. marinum*. Taken together, these observations indicate that a major curing event occurs early in the developmental program and likely at the single-cell level.

### Microfluidics to monitor *D. discoideum* at the single-cell level for several days

In order to decipher the cell-autonomous process of curing, we decided to dissect infection at the individual cell level. To that end, cells were trapped in a microfluidic device called the “InfectChip” (Fig S2 A) (Delince et al., 2016), especially suited to study long-term single-cell amoeba-bacteria interactions. The InfectChip is especially suitable for this task because of two reasons. First, *D. discoideum* cells are trapped into individual cages, which prevents them to physically interact and aggregate. Besides, the continuous perfusion of repletion/starvation medium in this device disrupts the cAMP gradient, preventing the cells to form streams. Since infection could impact the survival of *D. discoideum*, fitness in the InfectChip was assessed first in non-infected cells (Fig S2, Movie S2). To that end, the replication time and cell size were quantified over time. There was a slight lengthening of the replication time of *D. discoideum* cells in the InfectChip (14.01 h, Fig S2 B and Table S1) compared to the exponential growth in suspension culture (8.73 h, Fig S2 C and Table S1). This may be due to the 25 kDa cutoff of the semipermeable membrane, which might prevent cells from conditioning their medium by accumulating positive secreted factors that would continuously be perfused away. Alternatively, it might block the exit of negative factors that slow cell replication (Loomis, 2014). In any case, the replication time was more homogeneous in the InfectChip than in the various phases of growth in suspension culture. Indeed, during the first 12 hours of the lag phase, the replication time of *D. discoideum* is considerably longer (41.15 h, Fig S2 C and Table S1) than in the later exponential phase. We conclude that the fitness of the cells and their replication status in the InfectChip were suitable to study the infection.

We also assessed the fitness of *D. discoideum* in the InfectChip by measuring cell size during replication (Fig S2 D). Cells reached the same size before two consecutive divisions, thus implying cells obtain the necessary nutrients to increase their cell mass. In addition, only sporadic cell death was observed. As a conclusion, *D. discoideum* survived with the expected satisfactory fitness in the InfectChip for several days.

### Monitoring *M. marinum - D. discoideum* interactions using a microfluidic device

When *D. discoideum* cells infected with *M. marinum* by spinoculation were trapped in the InfectChip and imaged in repletion medium for several days, a surprising heterogeneity of events and infection fates was observed (Fig 3, Fig S3, Movie S3, Movie S4). First, infected *D. discoideum* cells continued to replicate and daughter cells inherited the bacteria in different fashions. We observed all kinds of distributions of bacteria during *D. discoideum* replication, from all bacteria being transferred to one of the siblings (Fig 3A), to a more homogenous distribution of the bacteria load among both daughter cells. It is to be noted that the former case would lead to a dilution of the number of infected cells, which could operate as a potential mechanism of curing at the population level. Regarding egress of the bacteria, *D. discoideum* released *M. marinum* via three mechanisms. During lytic release, *D. discoideum* cells died and bacteria frequently remained trapped within the debris (Fig 3B), facilitating re-uptake by bystander cells (Fig 3D). This process is important to spread the infection, and has already been described in human phagocytes (Hosseini et al., 2016; Mahamed et al., 2017). During non-lytic egress, *M. marinum* was released without compromising host viability (Fig 3C). As already described, non-lytic release comprises two distinct processes depending on the intracellular location of *M. marinum:* exocytosis of vacuolar bacteria, or ejection of cytosolic bacteria (Hagedorn et al., 2009) (Fig. 3C). These two processes were discriminated by monitoring actin. While LifeAct-GFP decorated the surface of the endosome prior to exocytosis (Fig S4 C), it formed a tight septum around the bacteria during ejection (Fig S4 D). The intracellular location of *M. marinum* (vacuolar or cytosolic) was also determined using specific endosomal markers that accumulate at the mycobacterium-containing vacuole (MCV), such as VacA-GFP (Fig S4 A, B, Movie S5). To our knowledge, this is the first time that the rupture of the compartment has been monitored with such a high spatial and temporal resolution. Similarly as upon lytic release, extracellular *M. marinum* released by exocytosis or ejection were frequently subjected to phagocytic reuptake by neighboring cells to start a new infection cycle. Sometimes, this phagocytic event occurred concomitantly to bacteria release, thus enabling the direct transfer of bacteria between hosts (Fig. 3E). Rarely, intracellular bacterial fluorescence disappeared, indicating killing and digestion of *M. marinum* (Fig 3F). All the events described above are summarized in the scheme presented in Fig 3G.

**Fig 3.**
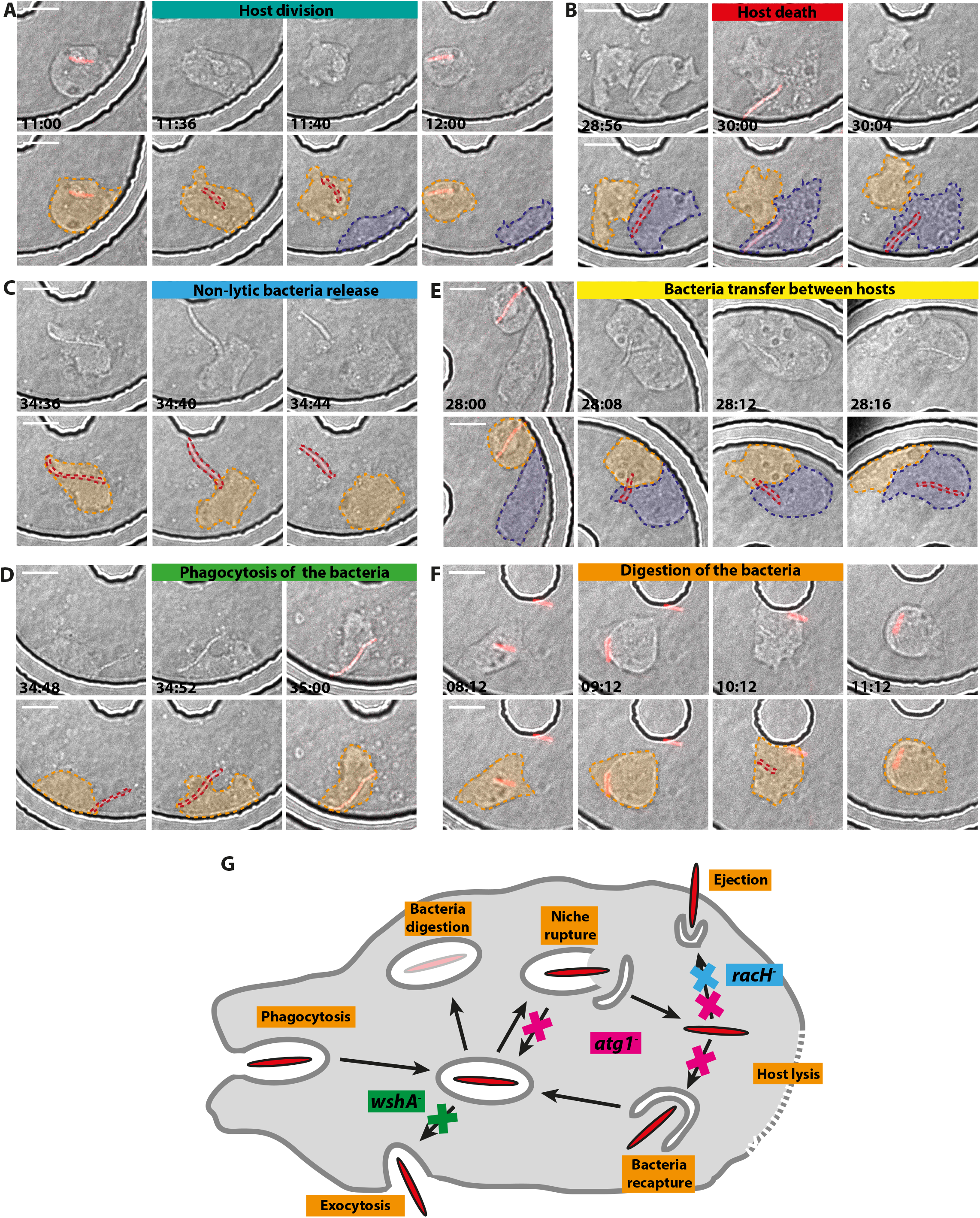
Classification of events observed during infection in the InfectChip microfluidic device. *D. discoideum* cells were infected with *M. marinum* wild-type or ΔRD1 (in red) and trapped in the InfectChip for live fluorescence microscopy. Six categories of events were recorded (upper panels merge of phase-contrast and fluorescence images, bottom panels pseudo-colored images). (A) Division of an infected cell gives rise to one infected and one non-infected cell. (B) Release of the bacteria by host lysis. (C) Release of the bacteria via a non-lytic mechanism (exocytosis or ejection, see Fig S4). (D) Phagocytic uptake of extracellular bacteria. (E) Direct bacteria transfer between a donor and an acceptor. (F) Bacteria fluorescence disappearance as a result of killing and digestion. (G) Schematic representation of the intracellular fates of *M. marinum* in *D. discoideum*. After phagocytic uptake, *M. marinum* transiently resides in a phagocytic compartment and can undergo different fates: exocytosis via fusion to the plasma membrane, killing and digestion of the bacteria via phagolysosomal maturation, vacuole escape via ESX-1-dependent membrane rupture. The autophagy pathway can prevent bacteria from escaping the compartment, either by sealing membrane damages or by engulfing cytosolic bacteria. Cytosolic bacteria can egress the host cell via ejection without lysing the cell. Finally, lytic death of the host at any stage can release the bacteria. Arrows and crosses indicate the steps impaired respectively in the mutants *wshA*-(exocytosis), *racH*-(ejection) and *atg1*-(resealing of the rupture compartment or recapture of the cytosolic bacteria in an autophagosome).

### The long intracellular residence time of *M. marinum* and subsequent death of *D. discoideum* are dependent on the RD1 locus

The “Region of Difference 1” (RD1) is the major pathogenicity locus described in *M. tuberculosis* and by homology, in *M. marinum* (Gao et al., 2004). To test the impact of this locus on the infection process, we used the InfectChip to quantify the bacterial fates previously described in *D. discoideum* cells infected with *M. marinum* wild-type or *M. marinum* ΔRD1. Mere inspection of trapped infected cells already suggested prominent differences between both bacterial strains regarding the length of the infection cycle and host viability (Fig S3 A, B, Movie S3, Movie S4). While wild-type bacteria seemed to undergo longer intracellular residency, ΔRD1 bacteria were released shortly after uptake, as depicted in the infection diagrams (Fig. 4A, E, Fig S5). Indeed, quantification revealed that 80.19% of ΔRD 1 bacteria were released non lytically in less than 10 h (Fig 4G, H and Table S2), compared to 45.10% when cells were infected with wild-type *M. marinum* (Fig 4C, D and Table S2). As a consequence, wildtype bacteria were able to grow intracellularly during this prolonged cycle (approximately 2-fold in the illustrative example of Fig 4B), whilst *M marinum* ΔRD1 barely showed a fluorescence increase (Fig 4F). As described before (Simeone et al., 2012), deletion of the RD1 locus also resulted in a vast reduction in host cell death, with 91.51% of the bacteria being released in a non-lytic manner (Fig 4G, H). In contrast, 27.45% of the *M. marinum* wild-type bacteria that remained intracellular for longer periods finally caused the death of their host (Fig 4C, D, Table S2). Taken all together, the mycobacterial RD1 locus seems to play an important role to establish a persistent infection, and it majorly impacts the viability of *D. discoideum*.

**Fig 4.**
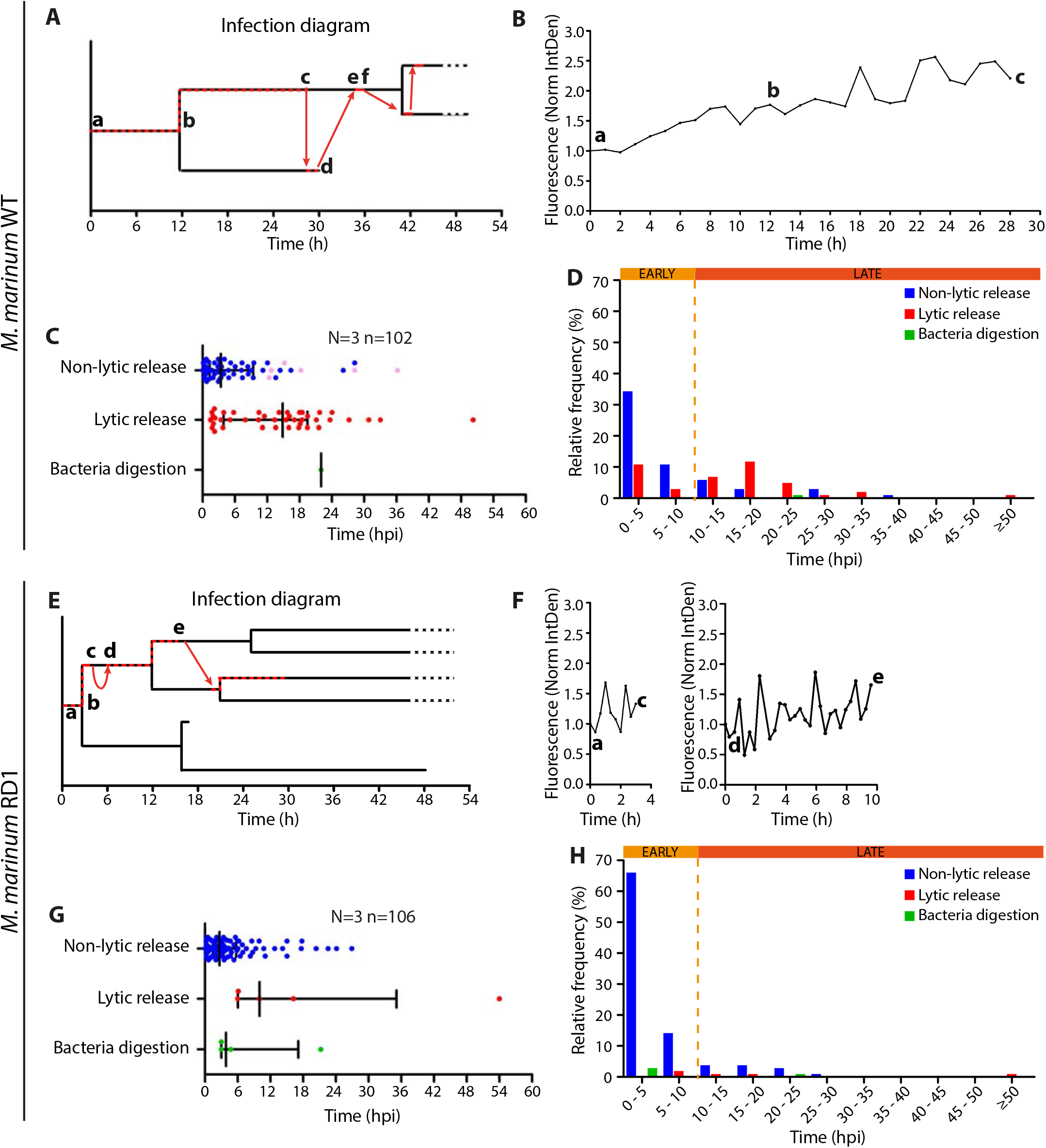
Dynamics of the infection of *D. discoideum* with *M. marinum* wild-type and ΔRD1. *D. discoideum* cells were infected with *M. marinum* wild-type or ΔRD1 and trapped in the InfectChip as for Fig 3 (see Movie S3 and S4 and Fig S3). (A, E) Fate diagram of the infections represented in Fig S3. Black lines trace the lineage of *D. discoideum* cells. Red dashed lines indicate presence of intracellular *M. marinum*. Red arrows represent release and reuptake of *M. marinum*. Letters refer to the events highlighted accordingly in Fig S3. (B, F) Intracellular growth of the bacteria during the intervals represented in B. (C, G) Scatter plots of the residence time of the bacteria before non-lytic bacteria release, lytic release or bacteria digestion. Median and quartiles are shown. The pink dots represent events that could not be followed until termination or that exceeded the duration of the recording. (D, H) Binned relative frequencies of the events represented in D.

### The intracellular fates of *M. marinum* are altered in the *D. discoideum* mutants *washA-, racH*- and *atg1-*

To further understand the intracellular fate and release mechanism of *M. marinum*, we decided to monitor in the InfectChip the course of infection in three *D. discoideum* mutants affected in major cellular pathways (*wshA-*, exocytosis, Movie S6; *racH-*, ejection, Movie S7; and *atg1-*, autophagy, Fig S5 D, Movie S8).

While wild-type *D. discoideum* cells constitutively exocytose undigestible remnants of pinocytosis and phagocytosis, *wshA*-knock out cells are greatly impaired (Carnell et al., 2011), if not completely blocked in postlysosome exocytosis. Correspondingly, when *wshA*-cells were infected with *M. marinum*, the early release of the bacteria was almost abolished (from 49.4% in wild-type cells down to 13.64% during the first 10 h, Fig S5 A, E, F, Table S2). As a consequence, the intracellular residence time of the bacteria increased substantially, with a median time for non-lytic release of 29.67 h compared to 3.40 h (wild-type), suggesting that this early process is dominated by exocytosis. In comparison, the median time for lytic release remained unaffected, with 13.73 h compared to 14.87 for wild-type (Fig 4C and Fig S5 E). Interestingly, this phenotype does not have an impact on the survival of the host (44.94% of events are attributed to the death of the infected *wshA*-cells, compared to 41.18.0% for wild-type cells, Fig S5 A, E, F, Table S2). In addition, bacteria were killed and digested slightly more frequently (6.74% in *wshA-*, 0.98% in the wild-type Fig S5 E, F, Table S2).

Host cells lacking the small GTPase RacH (*racH-*) do not form ejectosomes and accumulate cytosolic bacteria (Hagedorn et al., 2009). As expected, a decrease and delay of non-lytic release of bacteria was observed when *racH*-cells were infected and monitored in the InfectChip (16.07% during the first 10 h, Fig S5 B, G, H, Table S2). Similar to the infection in *wshA*-cells, increased bacteria retention (median of 25.67 h for non-lytic release or 21.33 h for lytic release, Fig S5 G) correlated with a loss of host viability (58.93% of bacteria causing the death of the host, Fig S5 B, G, H, Table S2). However, since *M. marinum* accumulates in the *racH*-cells at the cytosolic stage, it escapes the bactericidal endolysosomal pathway, reflected in a low bacteria killing and digestion, similar to the wild-type situation (Fig S5 G, H).

Finally, the autophagy pathway has been shown to play a complex and dynamic role at the interface with bacterial pathogens (Lopez-Jimenez et al., 2018). Consistent with this, in a mutant *D. discoideum* lacking the master autophagy regulator Atg1, *M. marinum* escapes to the cytosol prematurely, where it proliferates unrestricted by the xenophagic defence system (Cardenal-Munoz et al., 2017). In addition, autophagy has been shown to ensure plasma membrane integrity during ejection (Gerstenmaier et al., 2015). In summary, the load of both vacuolar and cytosolic *M. marinum* is controlled by autophagy, contributing to host cell defence against the infection. Despite this central role of autophagy, in the *atg1*-cells *M. marinum* did not incerase the frequency of host cell death (38.81% compared to 41.18% in wildtype cells, Fig S5 C, I, J, Table S2), which occured within the same time frame as in wild-type cells (median of 13.27 h for lytic release, compared to 14.87 h in wild-type cells, Fig S5 I, J). As might be expected, the frequency and median time for early release remained unchanged.

### Under starvation conditions, *M. marinum* is massively released from wild-type hosts in the first 10 hours

As a first step to decipher the reactions to starvation at the single-cell level, infected wild-type cells were starved inside the InfectChip. For that purpose, growth medium was perfused first and at 6 hpi. it was switched to starvation buffer (Fig 5A), in order to mimic the conditions in Fig 1A-D. A significant increase in non-lytic release events was observed (81.82%, Fig 5D, E, Table S2) compared to the control with growth medium (46.67%, Fig 5B, C Table S2). Furthermore, the non-lytic release of the bacteria occurred earlier (68.18% of the bacteria being released during the first 10 h post starvation onset, in contrast to 31.67% in repletion, Table S2). Concomitantly with the early release of the bacteria, there was a reduction in the frequency of host cell death under starvation conditions (17.05% compared to 46.67%, Table S2). This result confirmed that we can mimic the conditions leading to cell-intrinsic curing of *D. discoideum* in the InfectChip, and suggested that it mainly corresponds to an induction of non-lytic release of the bacteria via exocytosis or ejection.

**Figure 5.**
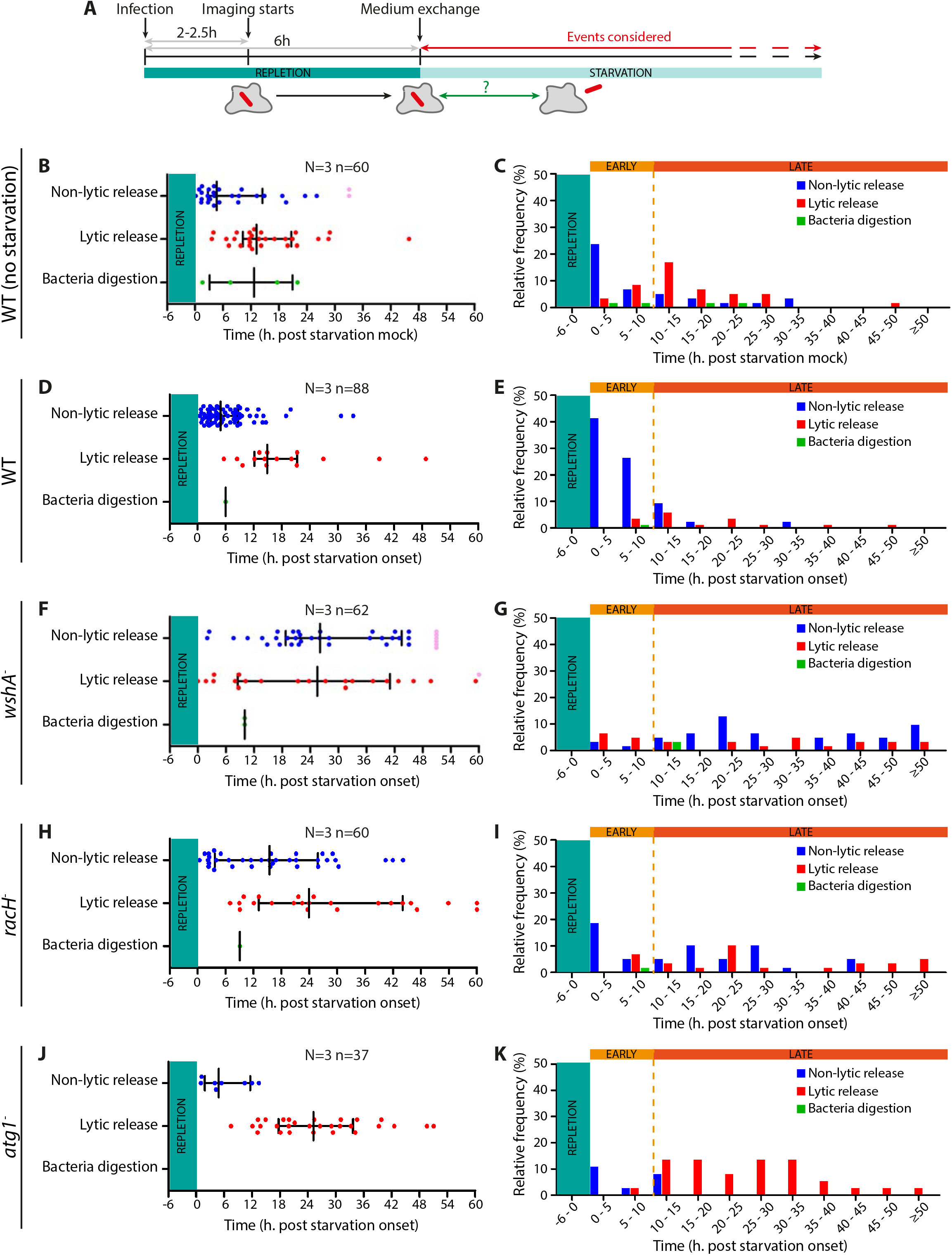
Early release of bacteria at the single-cell level during starvation does not occur in the *wshA-, racH*- and *atg1-D. discoideum* mutants. (A) Schematic representation of the experimental setting. At 6 hpi, perfusion in the microfluidic device was switched from repletion to starvation medium. This time, only the fate of bacteria that remained intracellular from the beginning of the imaging to the start of starvation were considered. Wild-type or mutant (*wshA-, racH*- or *atg1-*) *D. discoideum* cells were infected with *M. marinum* and trapped in the InfectChip for time-lapse live microscopy as described in Fig 3. *D. discoideum* cell lines and experimental conditions used are indicated in the left margin. (B, D, F, H, J) Scatter plots of the residence time of the bacteria before non-lytic bacteria release, lytic release or bacteria digestion. Median and quartiles are shown. The pink dots represent events that could not be followed until termination or that exceeded the duration of the recording. (C, E, G, I, K) Relative frequency of the events represented in B, D, F, H, J.

### The early release of *M. marinum* is impaired in the *wshA-, racH*- and *atg1*-mutants

Then, we decided to address which of the cellular pathways analysed in growth conditions (exocytosis, ejection or autophagy) contribute to cell-autonomous curing of the *M. marinum* infection induced by starvation.

The early release after starvation onset observed in wild-type cells (Fig 5D, E) was abolished in all three mutants (Fig 5F-K). In particular, non-lytic release of the bacteria during the first 10 h after the switch to starvation medium was reduced to a frequency of 4.84% in the *wshA*-(Fig 5G and Table S2), of 23.33% in the *racH*-(Fig 5I and Table 2), and of 13.51% in the *atg1*-(Fig 5K and Table 2), compared to 68.18% for wild-type cells under the same conditions (Fig 5E and Table S2). In addition, *M. marinum* seemed to follow similar fates in these mutant cells during starvation as quantified during continuous repletion (Fig S5). Indeed, infection in the *wshA*- and *racH*-cells were still characterized by delayed non-lytic release upon starvation (median of 26.07 h and 15.47 h respectively, Fig 5F, H), concomitant with an increase in host cell death, from 17.05 in wild-type cells (Fig 5E) to 35.48% and 38.34%, respectively (Fig 5G, I). Taken all together, the early release of *M. marinum* during *D. discoideum* starvation seems to be mediated by both exocytosis and ejection, and less intuitively also by autophagy.

### Curing at the collective level relies on exclusion of infected cells during aggregation

While 68.18% of intracellular *M. marinum* can be released via non-lytic mechanisms from their *D. discoideum* host during the first 10 h of starvation (Fig 5E and Table S2), 9.20% of cells remain infected even when starved for more than 20 h (Fig 5E). We thus reasoned that a second complementary mechanism of curing was intervening at later developmental stages in order to ensure uninfected aggregates and fruiting bodies.

Using live fluorescence microscopy, we monitored how elongated, polarized and streaming cells form aggregates and observed that rounder, infected cells were often found at the periphery of the tightly packed streams (Fig. 6A-G), or were even simply left behind. Such round infected cells are clearly observed at the periphery of streams when presented as serial xz (Fig. 6B-D) and xy-sections (Fig. 6F-G).

**Figure 6.**
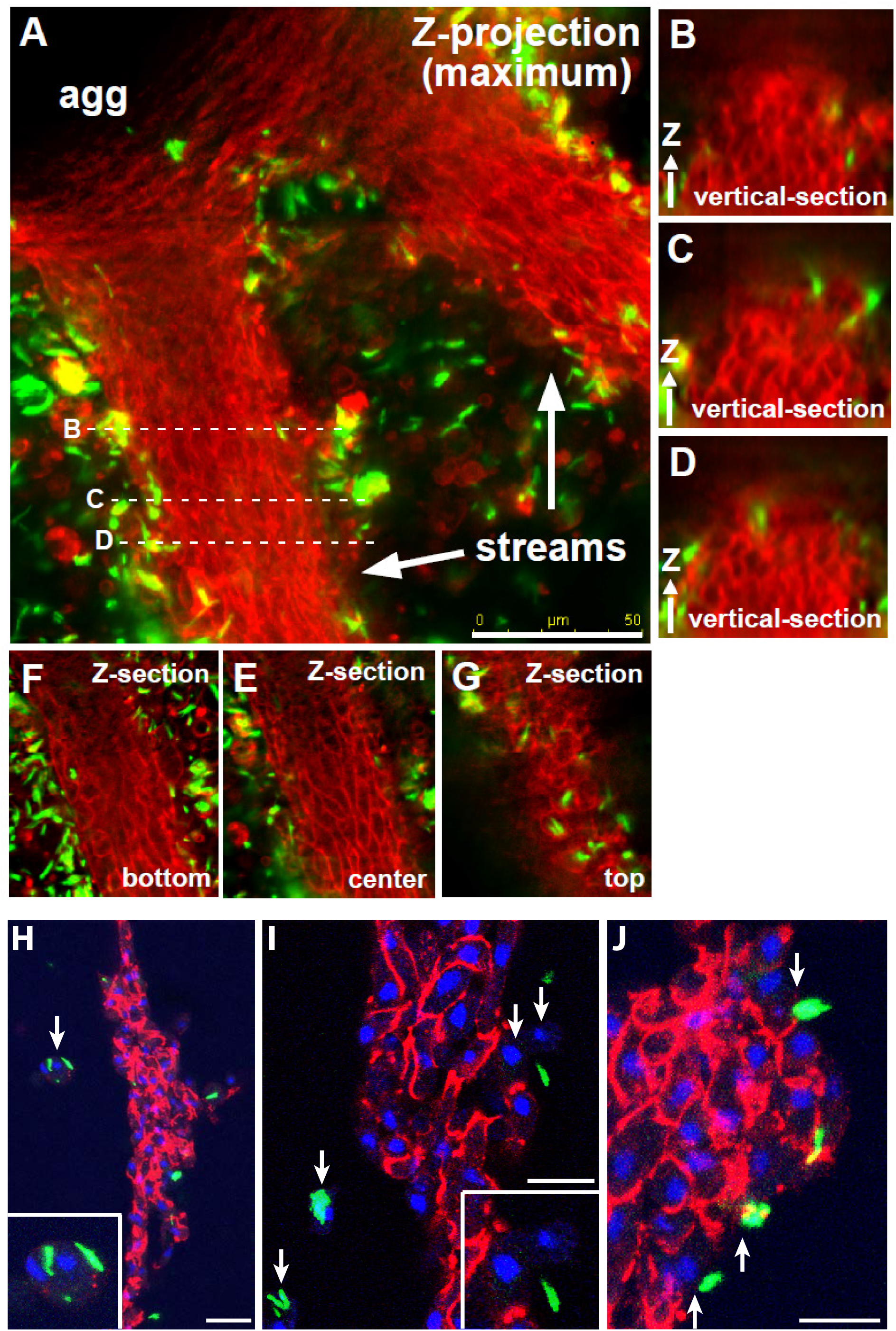
Infected *D. discoideum* cells are excluded from multicellular streams. (A-G) *D. discoideum* wild-type cells (red, stained with FM-464) were infected with *M. marinum* (green) and starved in Sorensen buffer from 12 hpi. An aggregate in formation (agg) with its feeding streams (arrows) was analysed by widefield live-cell fluorescence microscopy. (A) Maximum projection of 12 z-sections (9.6 μm) through the streams and aggregate. (B-D) Projections of vertical sections through the left stream. (F-G) single serial xy-sections from bottom to top of the stream. Positions of the z-sections in the streams are indicated in (A) with a dotted line. (H-J) Streaming cells were fixed and antibody-stained against contact site A (CsA). (H, I) CsA-expression and localization to the leading edge (red) of the polarized cells forming the stream (nuclei in blue). Infected cells are found at the periphery of the stream or completely excluded (arrows). (I) Higher magnification of infected cells shows that they do not express CsA and are not elongated. (H-J) Cells at the periphery of the stream that are associated with bacteria also show no signal for CsA, in contrast to their neighbouring streaming cells. Scale bars 50 μm (A) and 10 μm (H-J).

Streaming is a highly coordinated process during which cells express cell-cell adhesion molecules, such as Contact site A (CsA) (Siu et al., 2011). Immunofluorescence microscopy of cells at the streaming stage showed that, contrary to the cells forming the body of the streams and aggregates, the infected cells left behind and those found at the periphery of the streams did not express CsA (Figure 9H-J). This indicates that, lacking the appropriate cell-adhesion apparatus, infected cells are excluded from the streams, sorted towards the periphery, accumulate around the aggregate, and subsequently at the basal disk (Fig. 1G’, H’).

### *D. discoideum* mutants are impaired in sori sterility

To address the combined contributions of both the single-cell and the collective curing of the infection, we developed an assay to assess the sterility of the sori. For that, infected *D. discoideum* cells were allowed to aggregate on agar at 6 hpi and sori were collected after 2 days to measure bacteria content by CFUs (Fig. 7A). Even at a high MOI (30), the number of bacteria remaining associated with the sori was extremely low, thus indicating the extraordinary efficiency of the overall curing process (Fig 7B). As depicted in Fig. 7D, *racH*-sori harbored between 10.9 times more bacteria than wild-type. Surprisingly, *wshA*-cells seemed more capable to control the bacteria burden, with a similar number of mycobacteria CFUs remaining in the sori compared to wild-type (Fig 7C, D). Since both *racH*- and *wshA*-cells were severely impaired in the single-cell curing, this result may indicate a different ability of those mutants to overcome bacterial burden during the collective phase of development. Visualization of the sori of all these strains by fluorescence microscopy confirmed these results (Fig 7 E). Whereas wild-type sori remained almost uninfected, with only few bacteria mainly associated with the outside surface of the cellulose sheet, *wshA*-sori harboured small numbers of bacteria and *racH*-sori more severely contaminated with *M. marinum*. The sori-sterility assay could not be performed with the *atg1*-mutant because it does not reach the fruiting body stage of development. However, microscopy inspection of the loose aggregates formed by the *atg1*- at late stages contained high numbers of round infected cells.

**Fig 7.**
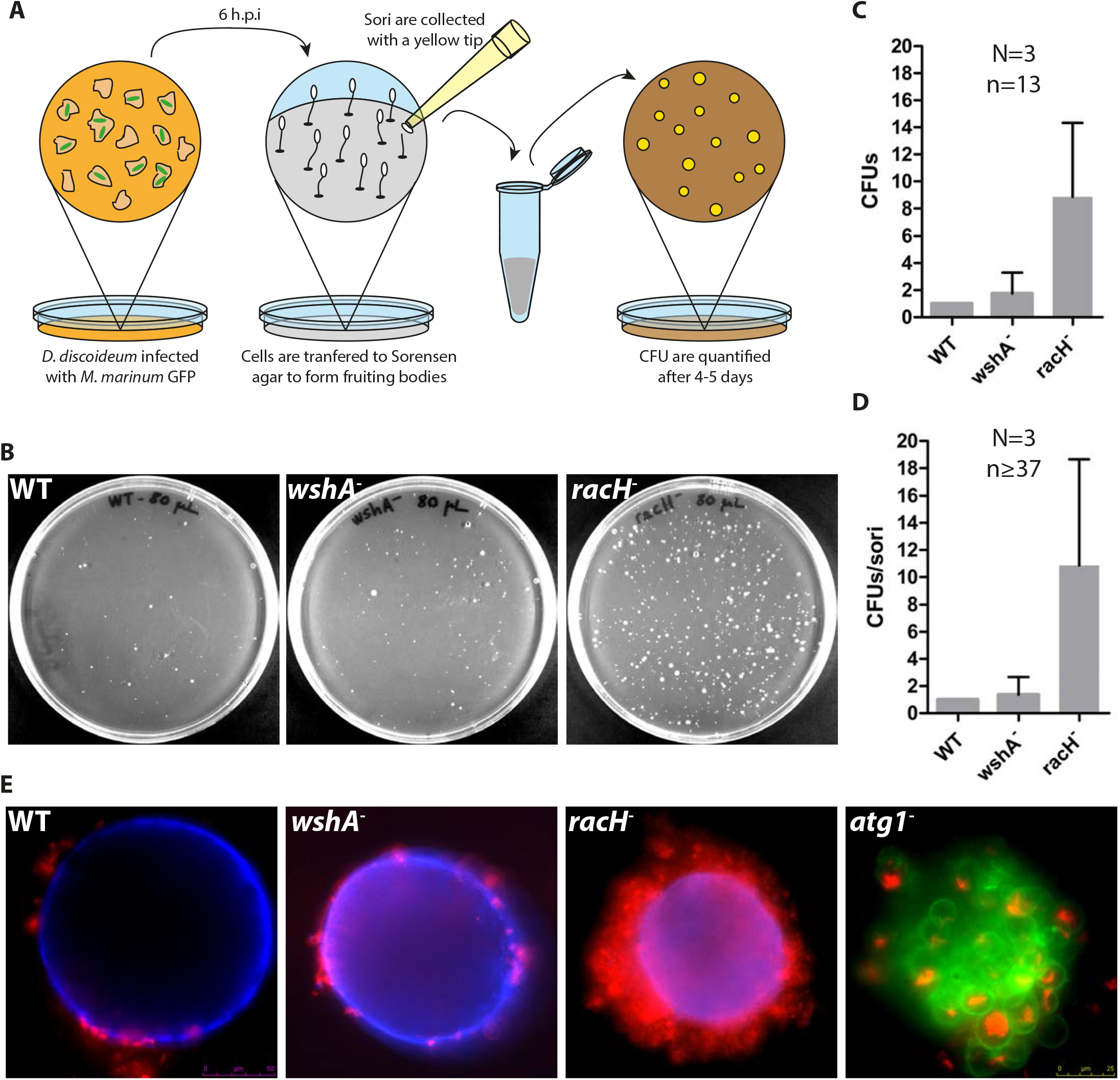
*racH*-is severely impaired in the formation of sterile sori. In comparison to wild-type *D. discoideum* (H), infected mutant strains with reduced capacity in ejection (*racH-*), exocytosis (*wshA-*) and autophagy (*atg1-*) show varying degrees of curing in the final aggregates. (A) Schematic representation of the experimental design to monitor sori sterility. Wild-type or mutant *D. discoideum* cells were infected and plated on non-nutrient-agar at 6 hpi. Bacteria presence in the sori was assessed by CFUs counting on 7H11 agar plates. (B) Representative 7H11 agar plates with *M. marinum* colonies for *wildtype, wshA*- and *racH*- (C) *M. marinum* CFUs of 13 sori from 3 independent experiments plated individually, normalized to wild-type. (D) *M. marinum* CFUs of 37 sori from 3 independent experiments for wild-type, *wshA*- and *racH-*. Sori from each experiment were plated together in this case. (E) Widefield images of sori from wild-type, *wshA*- and *racH*-(blue calcofluor, red *M. marinum*). The atg1-mutant has been shown to only reach the loose aggregate stage, which is also shown (red *M. marinum*, green *D. discoideum*, stained with FM-4-64). While the wild-type strain is very efficiently cured and shows no bacteria left in the cellulose-ensheathed aggregate, structures formed by the *racH, wshA*- and *atg1*-mutants still contain various numbers of infected cells.

## DISCUSSION

Although the first reports about the sterility of *D. discoideum* spores date more than 80 years ago (Raper, 1935), little progress has been made to understand the underlying mechanisms. Here, we analysed in detail the curing of a *D. discoideum* population highly infected with the pathogenic intracellular bacterium *M. marinum* using various microscopy and quantitative techniques. This enabled the identification of two phases in the curing process operating in a consecutive and complementary manner: the early cell-autonomous curing, consisting in the non-lytic release of bacteria; and the late collective curing, characterized by the collective exclusion of the remaining infected cells from the forming streams and aggregates (Fig 8).

**Figure 8.**
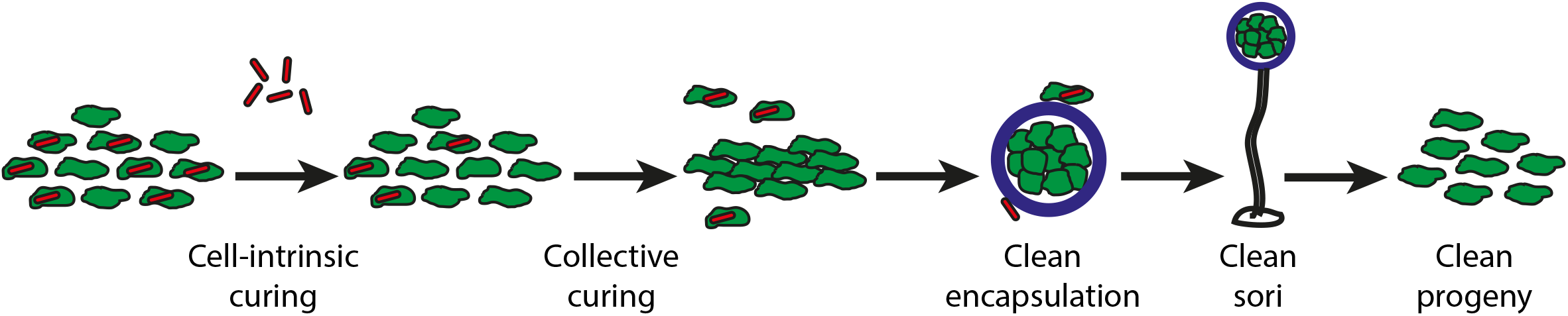
Model for curing of *M. marinum* infection upon starvation of *D. discoideum*. Under starvation conditions, infected *D. discoideum* cells (green) undergo two distinct phases of curing of *M. marinum* infection (red). In the early cell-autonomous curing phase, bacteria are released from single cells mainly via non-lytic mechanisms (exocytosis and ejection), and partly upon lytic host cell death. In the late collective phase, remaining infected cells are excluded from the forming aggregates since they do not express the adhesion molecule CsA. Sorting out and exclusion of infected cells lead to the encapsulation (cellulose sheath in blue) of noninfected cells in multicellular structures, the slug and then the sorus. Infected cells and bacteria stick on the outside and remain at the basal disc during fruiting body formation. This sequence of events results in uninfected sori and progeny.

Utilization of the “InfectChip” microfluidic device (Delince et al., 2016) was instrumental to understand the early cell-autonomous curing following starvation onset, and also revealed new insights in the infection process itself. The intracellular trafficking and main infection stages of mycobacteria in *D. discoideum* had already been extensively studied at the bulk population level (Cardenal-Munoz et al., 2017; Hagedorn et al., 2009; Hagedorn and Soldati, 2007). However, long-term single-cell imaging of infected cells in the InfectChip enabled a more exhaustive tracking and quantification of the events during infection and their interconnections. For instance, death of *D. discoideum* cells is a major fate and it was un underestimated in previous studies at the population level. There are two possible explanations for this. First, death happens within few minutes and cell corpses are fragile and usually lost during experimental handling. Second, death of *D. discoideum* cells is frequently followed by the partial or total phagocytosis of the cell remnants by neighbouring cells. Because of this, it was revealed that host death is a very efficient mechanism of propagation of the infection, since bacteria often remain within the dead cell debris, which facilitates their phagocytic uptake. This process is somehow reminiscent of efferocytosis, the engulfment of whole apoptotic cells by animal immune phagocytes (Green et al., 2016). However, efferocytosis requires caspase-mediated signaling that is absent in *D. discoideum*. Precise quantification of *D. discoideum* death during infection enabled us to confirm that the mycobacterium ESX-1 secretion machinery is an important factor involved in host cytolysis (Fig 4), as already reported (Gao et al., 2004). Moreover, since *M. marinum* grew during its intracellular cycle (Fig 4), phagocytic uptake of bacteria clumps by bystander cells lead to a higher cytotoxic burden that increased the frequency and accelerated host death, as previously described (Mahamed et al., 2017).

On the other hand, other types of events were difficult to monitor even using microfluidics. Since *M. marinum* form aggregates during intracellular replication, the mere measurement of the total fluorescence of bacteria clumps was not sufficient to discriminate for example growth arrest or a balance between bacteria growth and killing. Therefore, we believe that our measurements of bacteria digestion, which refer exclusively to the fluorescence disappearance of a bacteria or a whole bacterial clump, might be underestimated. This problem could be overcome in future studies by using mycobacterium septum markers such as Wag31-GFP in order to differentiate individual bacteria and their growth phase (Kang et al., 2008; Santi et al., 2013).

Besides, single-cell imaging revealed that infected cells are heterogeneous and follow very diverse fates. Indeed, microfluidics is emerging as a powerful tool to explore cellular individuality within a population and its role in shaping many biological processes, such as adaptation to changing environmental conditions, response to stress, and creates division of labour (Ackermann, 2015). Importantly, in the infection field, single-cell techniques have enabled the study of bacteria persistence to antibiotics, which relies on phenotypic heterogeneity (Dhar et al., 2016; Harms et al., 2016). During *M. marinum* infection in *D. discoideum*, this heterogeneity was reflected not only on the existence of multiple fates that intracellular bacteria could follow (summarized in Fig 3G), but also in the wide range of time spans in which these events occurred, which had already been reported (Delince et al., 2016). As an example, non-lytic release of *M. marinum* by *D. discoideum* happened after a few minutes up to several days (Fig 4).

The early cell-autonomous infection curing of *D. discoideum* seemed to rely predominantly on non-lytic bacteria release (Fig 5). This process is dependent on functional exocytosis, ejection and autophagy pathways, as demonstrated using the *wshA-, racH*- and *atg1*-mutants. During the early phase of infection, most bacteria remain in a modified phagocytic compartment and can undergo exocytosis upon starvation. Such a strategy is plausible, as it is known that the starvation process severely impacts the endocytic pathway (Smith et al., 2010) and differenciation induces exocytosis of phago-lysosomal enzymes (Smith et al., 2010). We hypothesize that prolonged intracellular residency might be responsible for increased bacterial toxicity and augmentation of *D. discoideum* death events. In addition, a small percentage of bacteria might be inherently less efficient at bypassing phagosomal maturation and consequently, the exocytic blockade in *wshA*-cells might favour their killing and digestion.

At later stages of infection, *M. marinum* translocates from its replication vacuole to the host cell cytosol, from which it is released through a plasma membrane ejectosome either to the extracellular medium or to neighbouring cells via concerted ejection-phagocytosis (Hagedorn et al., 2009). Therefore, induction of ejection might contribute to decrease the number of cytosolic bacteria. In *racH*-cells, which are defective in ejectosome formation, we did not observe any substantial increase in bacteria digestion, in contrast to the *wshA*-mutant. We believe that in *racH*-cells, *M. marinum* accumulates in the cytosol of its host, where it is not affected by the bactericidal activities of the phagocytic pathway.

Finally, starvation is the prime inducer of autophagy, and investigation of autophagy in the context of infection (xenophagy) has revealed its complex and dynamic role in controlling bacterial burden (Cardenal-Munoz et al., 2017). Both vacuolar and cytosolic *M. marinum* might be targeted by autophagy, captured and degraded, contributing to decrease the bacterial load and eventually cure host cells from infection.

The late collective curing process allows the formation of streams and aggregates exclusively formed by the non-infected *D. discoideum* population. Here, we show that cells that remain infected with *M. marinum* do not express the adhesion molecule CsA, required for the formation of compact pluricellular structures (Fig 6). However, the mechanisms that might control differential expression of CsA in developing cells depending on the presence of intracellular bacteria remain to be explored. Another plausible and untested scenario to explain the collective curing is that infected cells might also be unable to respond to the chemotactic cAMP signals.

Apart from these two major curing processes, S-cells in the slug have also been shown to have bactericidal functions, either by phagocytic killing or by casting DNA extracellular traps (Zhang et al., 2016). Besides, it should not be excluded that residual infected cells in the slug may differentiate preferentially into stalk cells rather than becoming spores. These later stages of the curing process might contribute to a better control of the bacteria burden in the sori by the *wshA-cells*, whereas the *racH*-cells might be impaired.

Overall, as a result of the multilayered curing mechanisms, wild-type *D. discoideum* cells are extremely efficient at generating sori devoid of pathogenic *M. marinum*. The obvious biological consequence is the formation of sterile spores and an uninfected progeny of vegetative cells, which may confer an ecological advantage.

## MATERIALS AND METHODS

### *D. discoideum* cell culture

Wild-type *D. discoideum* (Ax2 and Ax2(Ka)) and mutant cell lines were cultured axenically at 22°C in HL5c medium (Formedium). The *D. discoideum* mutant Ax2G *wshA*-was kindly provided by Dr. R. Insall (King et al., 2013), Ax2G *racH*- by Dr. F. Rivero (Somesh et al., 2006), Ax2(Ka) *atg1*- by Dr. J. King (King et al., 2013) and DH1 *atg1*-(DBS0236336) was obtained from dictyBase (dictyBase.org). VacA-GFP- and GFP-LifeAct-expressing Ax2(Ka) *D. discoideum* were obtained by transformation with pDM1045 and pDM1043 vector respectively (Veltman et al., 2009). ABD-GFP was used before (Lee and Knecht, 2002).

### Mycobacteria strains, culture and plasmids

Wild-type *M. marinum* (M strain) and ΔRD1 were kindly provided by Dr. L. Ramakhrisnan (Cardenal-Munoz et al., 2017). Mycobacteria were cultured in 7H9 (Difco) supplemented with glycerol 0.2% (v/v) (Biosciences), Tween 20 0.05% (v/v) (Sigma) and OADC 10% (v/v) (Middlebrock) as described (Kolonko et al., 2014). GFP-, mCherry- and DsRed-expressing bacteria were obtained by transformation with msp12::GFP, pCherry10 and pR2Hyg respectively, which were kindly obtained from Dr. L. Ramakrishnan, (Carroll et al., 2010; Cosma et al., 2004; Cosma et al., 2008). Cultivation was performed in the presence of 25 μg/ml kanamycin (Sigma) for GFP-expressing bacteria and 50 μg/ml hygromycin (Labforce) for mCherry10 and DsRed-expressing *M. marinum*.

### Measurement of *D. discoideum* replication time in suspension

10^5^ *D. discoideum* cells were collected in a total volume of 10 mL of fresh Hl5c (Formedium). Cell density was measured every 12 h using a Scepter Automated Cell Counter (Merck). Replication time was calculated as

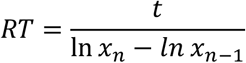

with *RT* the replication time, *t* the time and *x_n_* the number of cells at a certain time-point.

### Infection assay

Infections were performed as previously described (Arafah et al., 2013; Hagedorn et al., 2009). In brief, fluorescent mycobacteria were centrifuged (twice 500 g for 10 min) onto adherent *D. discoideum* cells. After an additional 20-30 min of incubation, free bacteria were washed off and the infected cells were resuspended in Hl5c containing 5 U/mL of penicillin and 5 μg/ml streptomycin (Thermo Fisher Scientific). The time of addition of bacteria to the adherent cells is referred to as 0 hours post infection (hpi). Different multiplicities of infection (MOI) were used depending on the subsequent assay performed. For the sori sterility assay, an MOI of 30 (Fig 7A-D) was used to reach a fraction of infected cells around 60-70%, for other differentiation assays the MOI was 100-300 (Fig 1 and 7E), so that a fraction of at least 80% was reached. For single-cell analyses in the InfectChip, an MOI of 10 was used with Ax2G, Ax2(Ka) and *wshA-*, and an MOI of 2 for *atg1*- and *racH-*. The percentage of infected cells was determined by flow cytometry or fluorescence microscopy.

### Differentiation assay

To initiate the differentiation process, *D. discoideum* growth medium (Hl5c) was removed, infected cells were washed twice with Soerensen buffer (SB, 14.7 mM KH2PO4, 2.5 mM NaHPO4, pH 6.3) before being allowed to differentiate in SB undisturbed in a humid chamber at 25°C.

To monitor the fate of fluorescent bacteria and differentiation to the slug-stage, live infected cells were seeded in an untreated 35 mm μ-dish (Ibidi) and covered with a thin layer of Sorensen buffer. The cells were imaged using a Leica LF6000LX microscope (100x or 63x objective, oil immersion) up to 28 hours at 25°C. Micrographs, brightfield and fluorescence (GFP and DsRed), were recorded every 5-15 minutes. To obtain a larger field of view the Leica-software (LAS) was used to stitch 4 neighbouring images together.

Full differentiation to mature fruiting bodies was achieved after plating the infected cells on 10 cm Sorensen-agar plates, which were monitored at different times using a binocular microscope (Nikon SMZ1000) equipped with a fluorescence lamp to excite GFP (excitation 488nm, emission 510 nm) and a Nikon Digital DXM1200F camera.

### Sori sterility assay

Up to 10^7^ *D. discoideum* cells infected with GFP-expressing *M. marinum* were washed once with Sorensen buffer and plated on 6-cm Sorensen-agar plates at 6 hpi. After two days, presence of fruiting bodies was assessed using a binocular microscope. Sori were picked with thin pipette tips, suspended in 100 μl of Sorensen buffer and plated on 7H11 agar plates supplemented with 2.5 mg/L Amphotericin and 25 μg/μL of kanamycin (Sigma). After 5 days of incubation at 32 °C, *M. marinum* colony forming units (CFUs) were counted.

### Immunofluorescence and stainings

The monocolonal antibody against CsA (41.71.21) was obtained from Dr. G. Gerisch. The secondary antibody was a goat anti-mouse IgG coupled to Alexa 488 (Life Technologies).

To stain the cellulose sheath surrounding aggregates, calcofluor white stain (FLUKA) was used at 0.5 mg/ml. The lipophilic dye FM4-64 (Life Technologies) was used to image live cells in streams according to manufacturer’s instructions.

### Imaging of streams and aggregates

To image streams stained for CsA, infected cells were allowed to stream in 35 mm μ-dishes (Ibidi). Subsequently, the streams were fixed with 4% PFA in SB for 1 hour, permeabilized and stained for CsA as described in (Hagedorn and Soldati, 2007). The stained cells were embedded using ProlongGold antifade (Life Technologies) and covered with a coverslip in the μ-dish. Imaging was performed with a Leica LF6000LX microscope (100x or 63x objective, oil immersion) and stacks were recorded. A blind deconvolution algorithm (LAS-software) was applied to deblur the image.

Aggregates of live cells were stained with calcofluor and FM 4-64 and imaged in a layer of SB without further fixation or embedding. Using a Leica LF6000LX microscope or Olympus IX81 microscope (100x objective, oil immersion; 1,4 NA), z-stacks (distance between sections of 0.4 to 1.8 μm) were taken through the aggregate and deconvoluted using either the LAS-software (blind deconvolution) or the nearest-neighbour algorithm integrated in the Xcellence rt software. Brightness and contrast of images were adjusted using ImageJ (http://rsb.info.nih.gov/ij/). The reslice and grouped maximum projection function in ImageJ was used to generate vertical and serial xy-sections through the streams of cells.

### Single-cell imaging in the InfectChip

The components of the InfectChip were kindly provided as a result of the collaboration with J.D. McKinney (EPFL, Lausanne, Switzerland) and assembled as described (Delince et al., 2016). Briefly, *D. discoideum* cells were seeded for 15-20 min on a SU8 GM1060 coverslip micropatterned using photolithography techniques. This coverslip is placed on a stainless steel bottom holder and covered with a semi-permeable membrane (25 KDa molecular cutoff pre-treated Spectra/Por R7 dialysis tubing, Spectrum Labs) that was washed, cut, dehydrated and rehydrated as previously described. On top of the semi-permeable membrane, a microfluidic chip of polydimethylsiloxane (PDMS) is placed. The PDMS microfluidic chip presents a serpentine channel and connected to two silicon tubes (0.76 mm inner diameter) for continuous medium perfusion. Finally, this device is held in place by screws tightly uniting the bottom holder with a PMMA upper holder. The tubes were connected to a 50 mL BD luer lock syringe loaded with the perfusion medium into the microfluidic device, such as Hl5c containing 5 U/mL of penicillin and 5 μg/ml streptomycin (Thermo Fisher Scientific) for repletion or Hl5c 5% (v/v) in Sorensen buffer supplemented with 100 μM of CaCl2 and 5 U/mL of penicillin and 5 μg/ml streptomycin for starvation. A programmable Aladdin Syringe pump was used to apply a constant flow of 10 *μ*l/min from the syringe.

*D. discoideum* cells trapped in the InfectChip were imaged at a Spinning Disc Confocal Microscope (Intelligent Imaging Innovations Marianas SDC mounted on an inverted microscope (Leica DMIRE2)) with an oil 100x objective. Z-stacks of 6 slices of 1.5 μm were acquired every 4 min in phase-contrast to track the cells, and every 20 min for *M. marinum* ΔRD1 or 1h for *M. marinum* wild-type in fluorescence in order to minimize phototoxicity.

### Image analysis for InfectChip experiments

The interdivision time was defined as the time between the first and the second observed cell division of a trapped cell. The projected surface occupied by a cell was measured in ImageJ after manual selection of the cell contour visible in phase contrast. Bacteria growth was calculated according to the following steps: (1) bacteria were identified by the creation of a mask using the red fluorescence channel as described in (Barisch et al., 2015); (2) the two slices in the z-stack with higher fluorescence were determined; (3) the integrated density of the selected slices was added. Vac-A GFP fluorescence was measured by creating a band-shape zone around a mask created using the fluorescence of the bacteria.

## ACKNOWLEDGMENTS

We gratefully acknowledge the members of the “Amoeba Club”, including Dr. M. Blokesch, Dr. P. Cosson, Dr. G. Greub and their groups for fruitful discussion. We thank the Bioimaging Center for Microscopy and Image Analysis at the Faculty of Sciences in Geneva for their precious help. The T.S. laboratory is supported by multiple grants from the Swiss National Science Foundation and T.S. is a member of iGE3 (www.ige3.unige.ch).

## SUPPLEMENTARY FIGURES

**Supplementary Figure 1.**
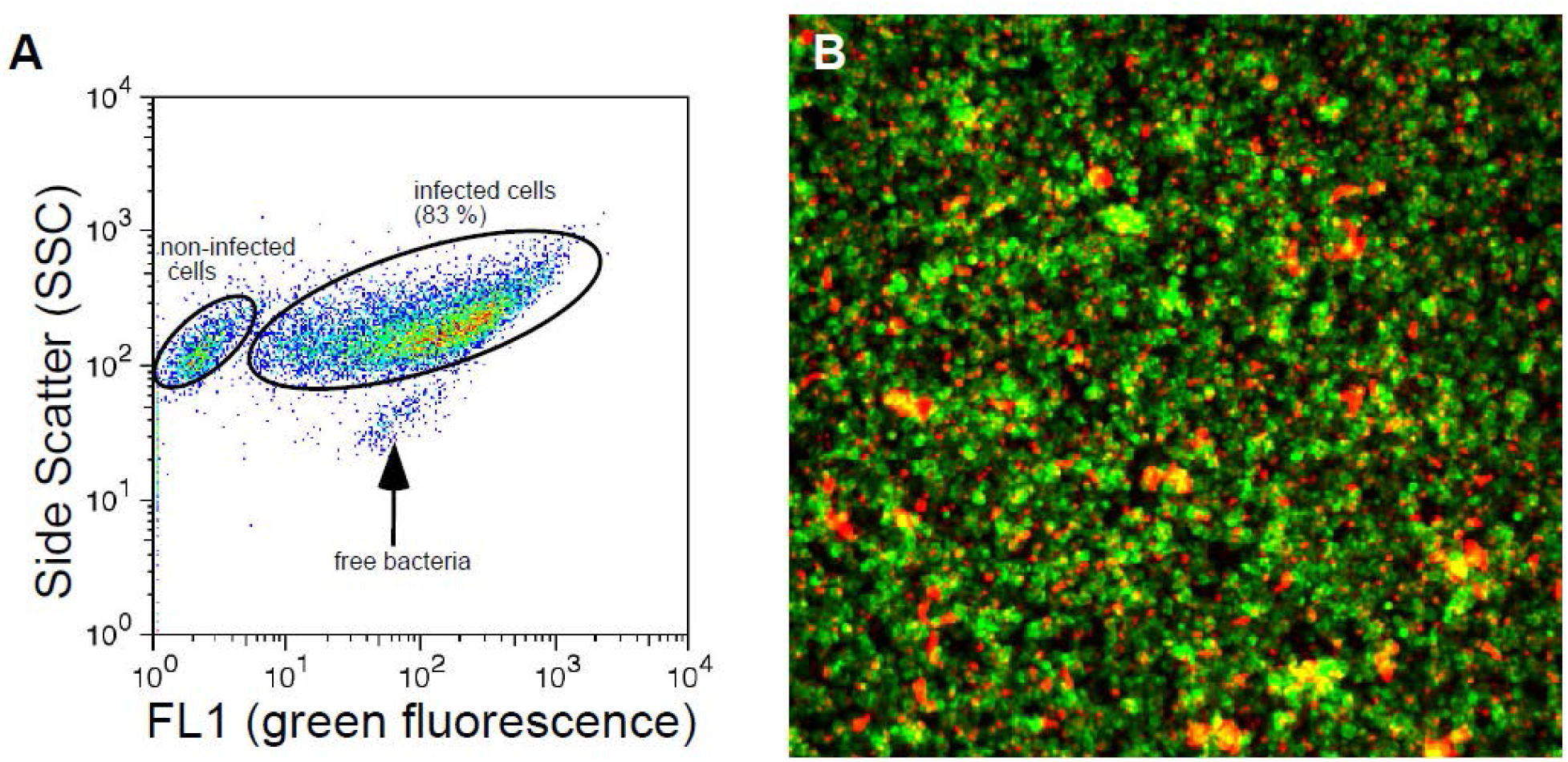
Efficient infection in *D. discoideum* cells. Efficient infection of *D. discoideum* cells was confirmed by flow cytometry (A) and fluorescence microscopy analyses (B). (A) *D. discoideum* cells were infected with GFP-expressing *M. marinum* and analysed by flow cytometry, and the green fluorescence (FL1, x-axis) plotted versus the side scatter (SSC, y-axis). Over 80% of the cells were infected with green fluorescent *M. marinum* and only very few free bacteria were observed. (B) Fluorescence microscopy was used to judge the infection efficiency and exclude the presence of large numbers of free bacteria. Close to 100% of green fluorescent *D. discoideum* cells carried red-fluorescent bacteria.

**Supplementary Figure 2.**
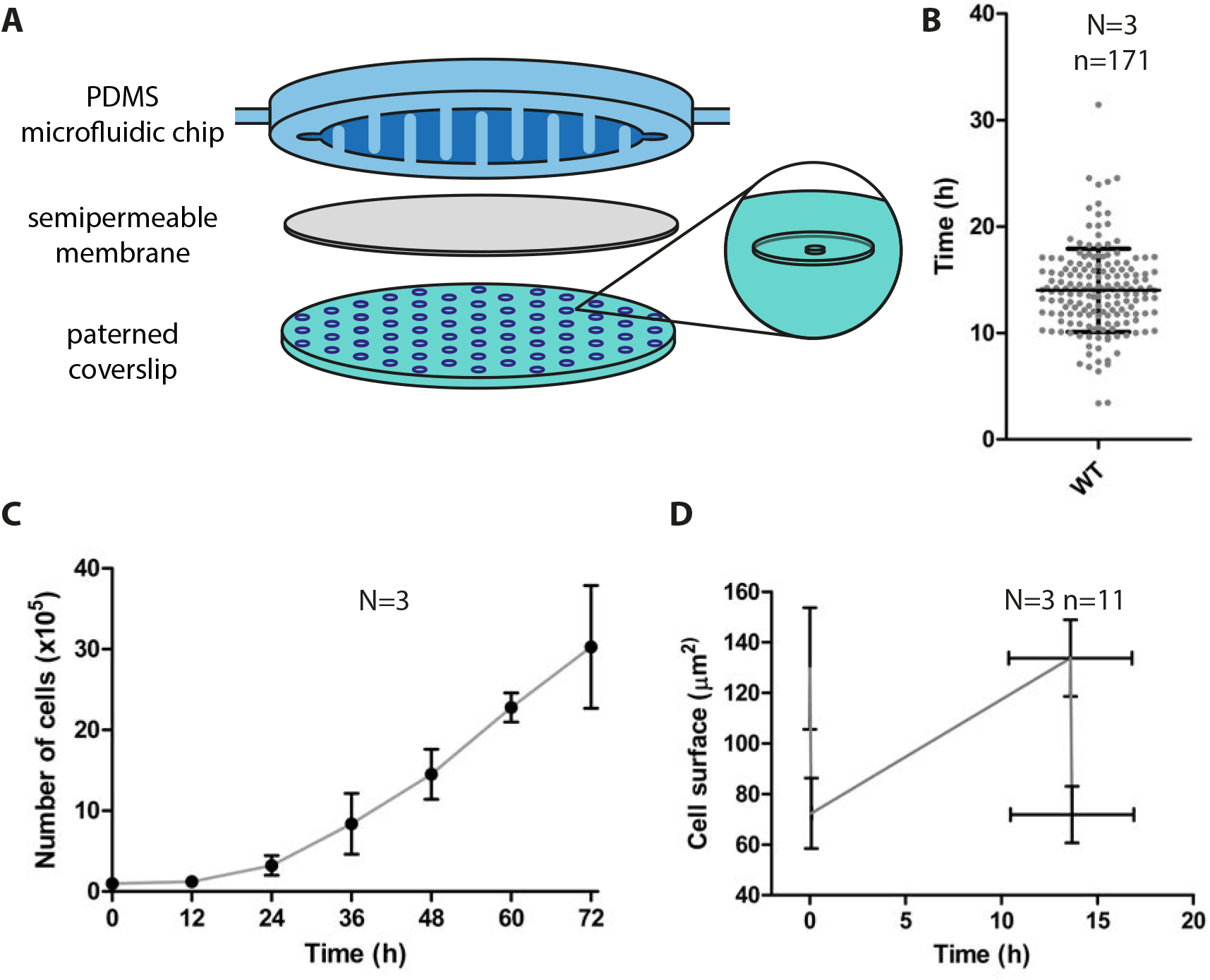
Fitness of *D. discoideum* cells in the InfectChip compared to growth in suspension. (A) Schematic representation of the InfectChip. (B, D) Non infected *D. discoideum* cells were trapped in the InfectChip and imaged at a spinning disc confocal microscope (phase-contrast images every 4 min, Movie S2) (B) Interdivision time of 171 consecutive cell divisions from 3 independent experiments. Graph shows mean and standard deviation. (C) Non infected *D. discoideum* growth in suspension in a conical flask. The graph shows the mean and standard deviation from 3 independent experiments. (C) Conservation of size by noninfected *D. discoideum* cells during consecutive divisions in the InfectChip. The graph shows the mean and standard deviation of the cell surface and time of replication of 11 cells from 3 independent experiments.

**Supplementary Figure 3.**
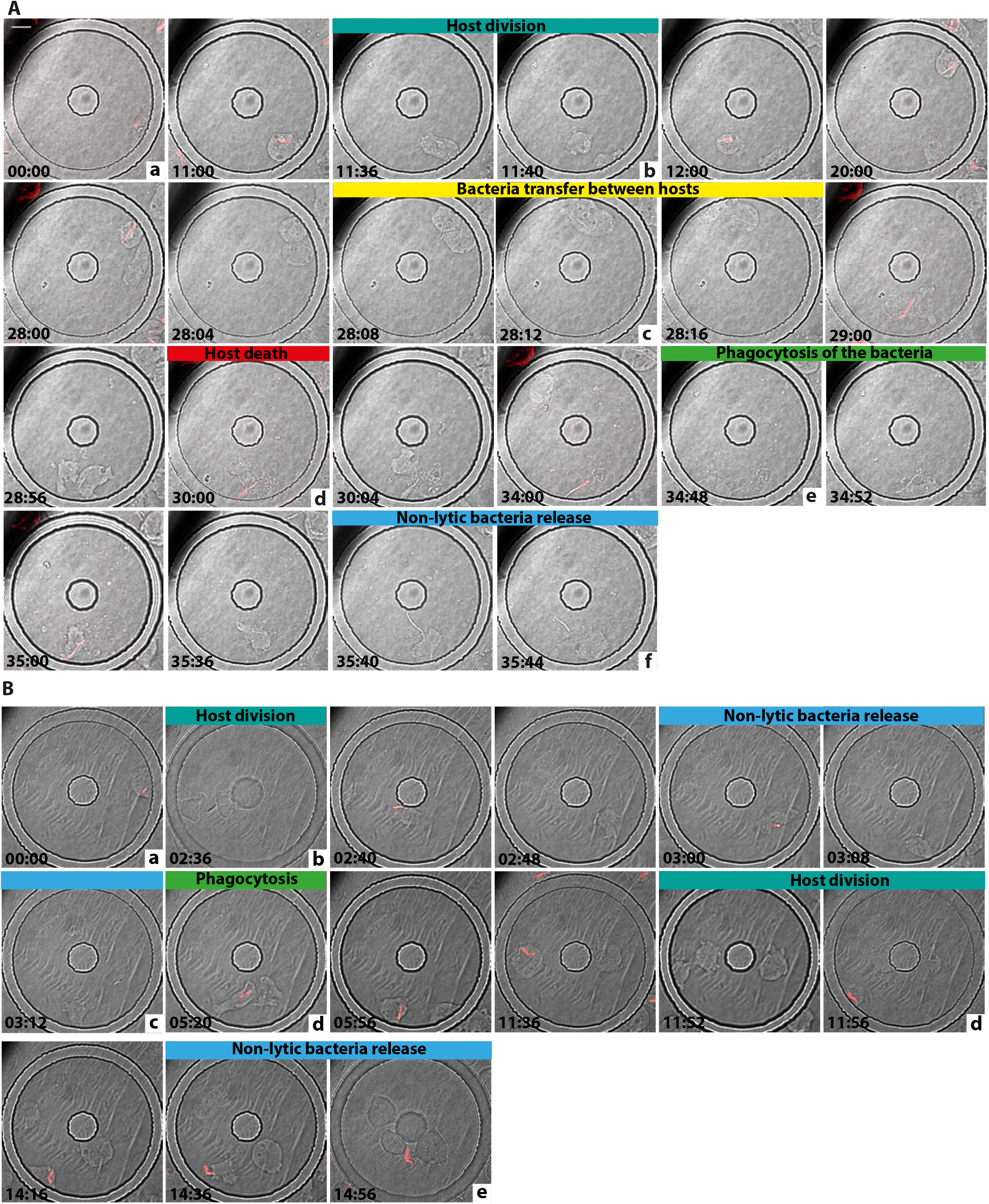
Infection of *M. marinum* and *M. marinum* ΔRD1 in trapped *D. discoideum* cells. Still frames from a 40h movie (Movie S3 and Movie S4) of *D. discoideum* infected with *M. marinum* (A) or *M. marinum* ΔRD1 (B). Letters correspond to events represented accordingly in Fig 4.

**Supplementary Figure 4.**
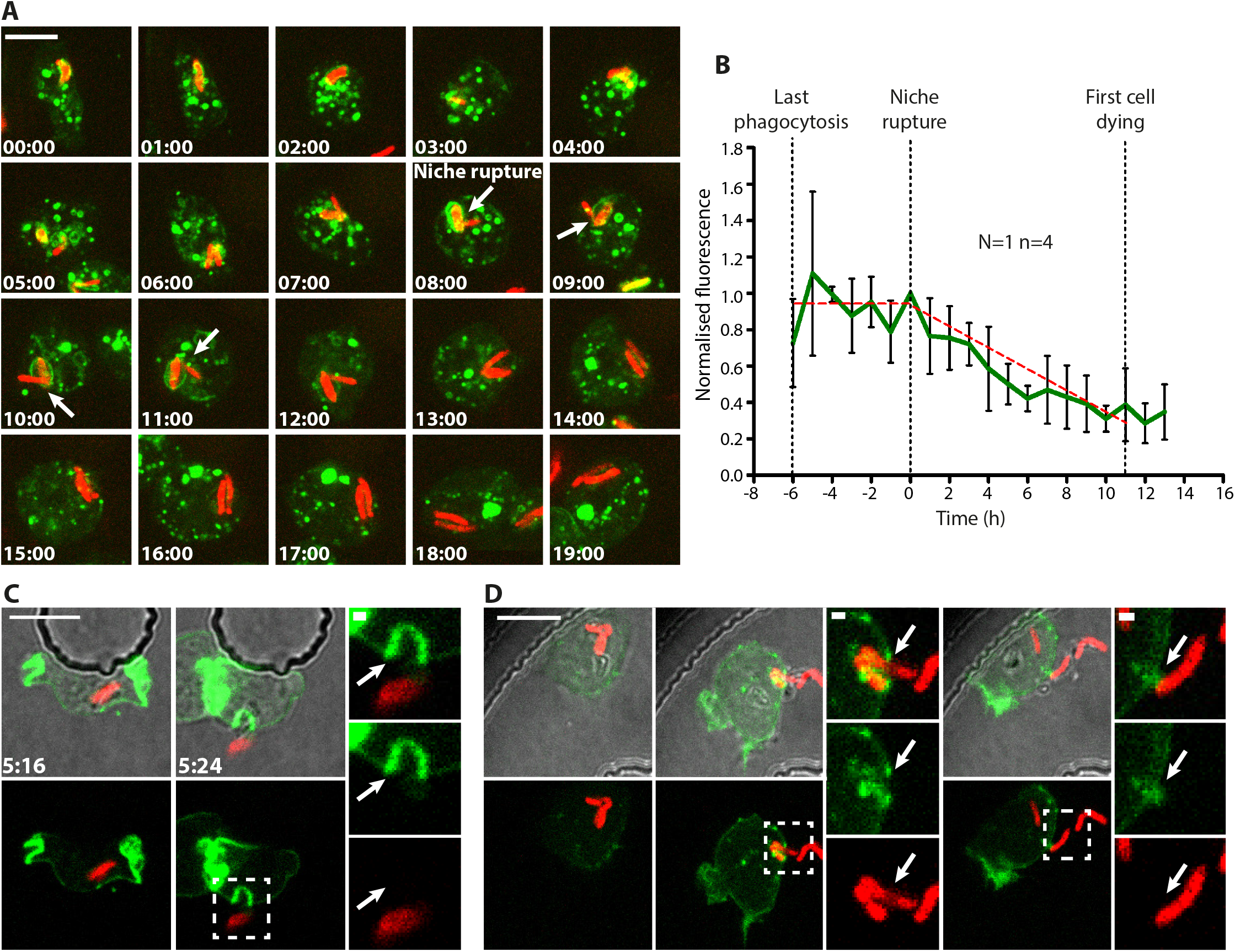
Niche rupture, exocytosis and ejection can be visualized using specific markers. (A) Localisation of GFP-VacA in *D. discoideum* enables precise monitoring of the MCV (bacteria in red). Time-lapse showing increase in GFP-VacA fluorescence at the MCV, rupture of the compartment (arrows) and disappearance of the fluorescence. (B) Quantification of fluorescence around the bacteria, normalized and temporally synchronized to the moment of niche rupture. GFP-VacA decorates the *M. marinum* containing compartment, until the moment of rupture, when fluorescence starts to disappear. (C-D) Localisation of LifeAct-GFP in *D. discoideum* enables the discrimination between exocytosis and ejection of *M. marinum*. (C) A LifeAct-GFP decorated MCV fuses with the plasma membrane during exocytosis of *M. marinum*. (D) A tight septum of fluorescent LifeAct forms around the bacteria being ejected. Insets show detail in the ejection process.

**Supplementary Figure 5.**
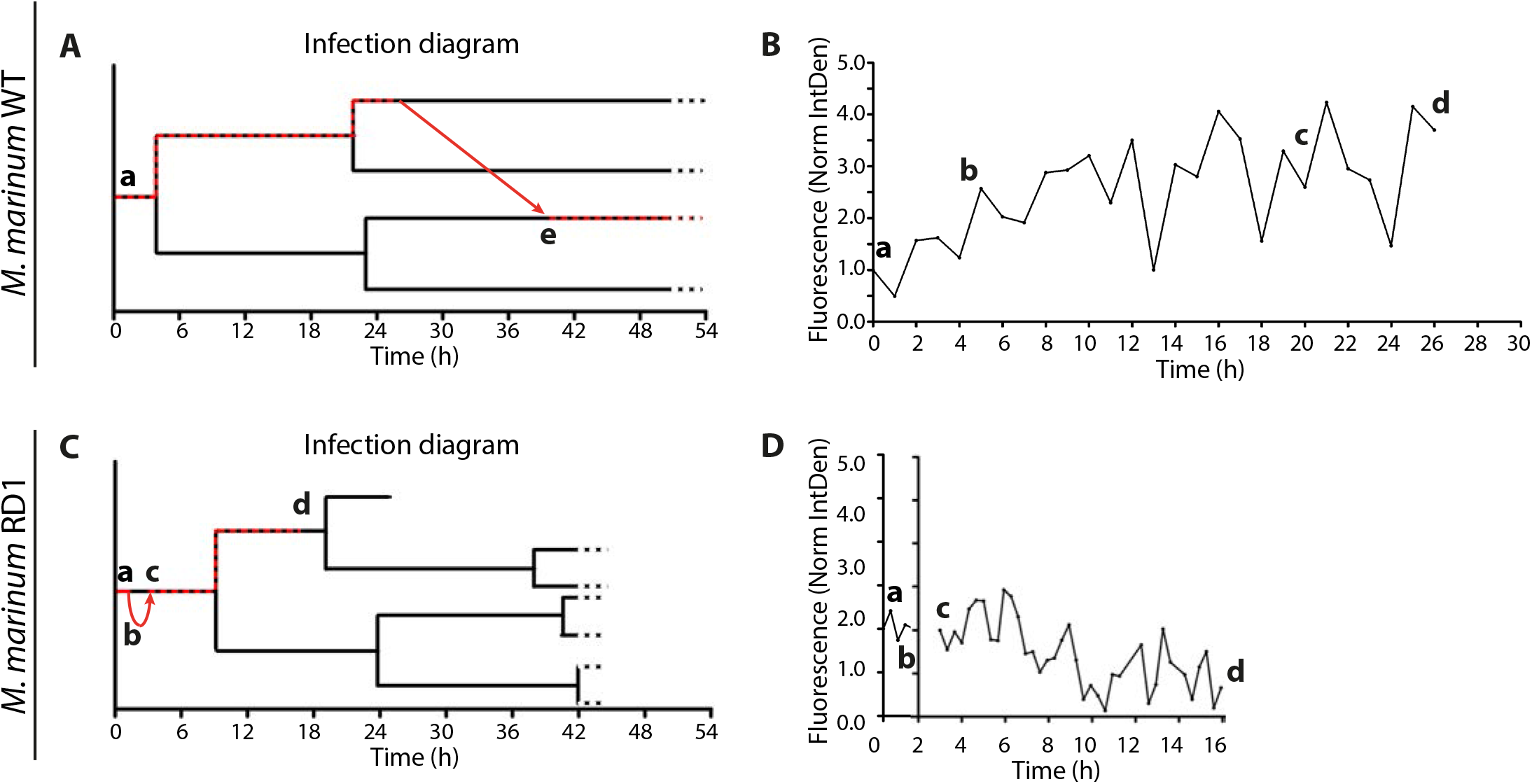
Differences in the infection cycle determined by the mycobacterial RD1 locus. *D. discoideum* cells were infected with *M. marinum* wild-type or ΔRD1 and trapped in the InfectChip as for Fig 4. (A, C) Additional fate diagram of representative infections with wild-type or ΔRD1 *M. marinum*. Black lines trace the lineage of *D. discoideum* cells. Red dashed lines indicate presence of intracellular *M. marinum*. Red arrows represent release and reuptake of *M. marinum*. Letters refer to the events highlighted (B, D) Intracellular growth of the bacteria during the intervals represented in A, B.

**Supplementary Figure 6.**
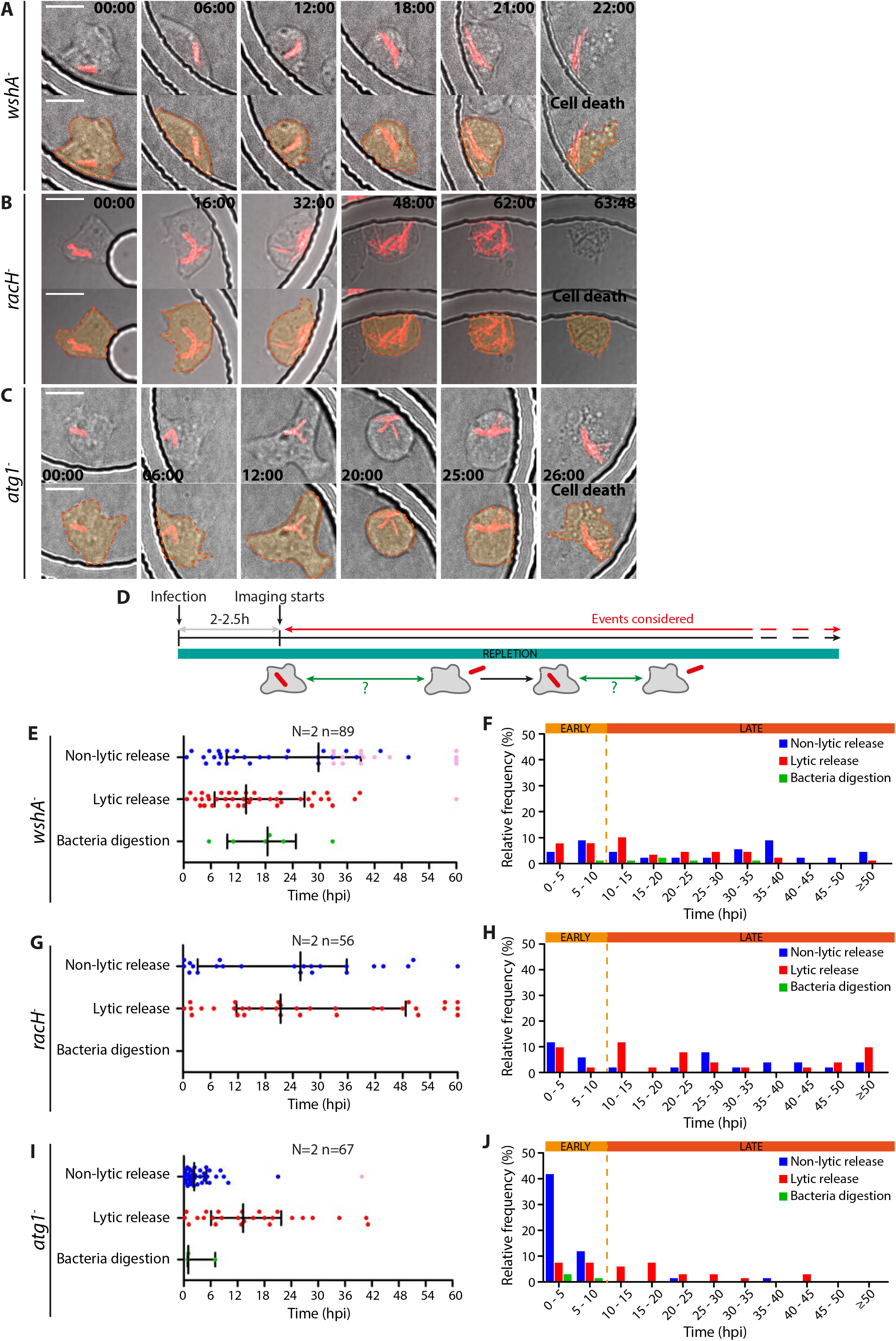
The fate of *M. marinum* is altered in *D. discoideum* lacking WshA, RacH and Atg1. The fate of *M. marinum* is impacted by the mutations *wshA-, racH*- and *atg1-*. (A-C) Still frames of 60 h movies of *wshA-, racH*- and *atg1-D. discoideum* was infected with *M. marinum* and trapped in the InfectChip as described in Fig 3. (D) Schematic representation of the experimental setting. (E, G, I) Scatter plot of the residence time of the bacteria before non-lytic bacteria release, lytic release or bacteria digestion. Median and quartiles are shown. In black are represented events that could not be followed until termination or that exceeded the labelling of the X-axis. (F, H, J) Relative frequency of the events represented in (E, G, I).

**Supplementary Table 1.** Summary table of *D. discoideum* growth in the InfectChip and in shaking conditions. Times of replication in the InfectChip and in shaking conditions, data complementary to Figure S2.

| InfectChip |  |  |  |
| --- | --- | --- | --- |
| Number of events | Mean (h) | Median (h) | StDv |
| 171 | 14.01 | 14.05 | 3.907 |
| Shaking suspension |  |  |  |
| Culture phase | Time | Replication time (h) |  |
| Lagging | 0h - 12h | 41.15 |  |
| Exponential | 12h - 24h | 8.73 |  |

**Supplementary Table 2.** Summary table of the bacteria fates during infection. Percentage of bacteria or bacteria aggregates following the mentioned fates during infection. “All events considered” and “Reuptakes excluded” refer to events quantified as represented in F S5 D or in Fig 5A, respectively.

| All events considered |  |  |  |  |  |
| --- | --- | --- | --- | --- | --- |
| *D. discoideum* | *M. marinum* | Condition | Early events (up to 10 h.p.i) (%) |  |  |
|  |  |  | Non-lytic release | Lytic release | Bacteria digestion |
| wild-type | wild-type | repletion | 45.10 | 13.73 | 0 |
| wild-type | ΔRD1 | repletion | 80.19 | 1.89 | 2.83 |
| *wshA*<sup>-</sup> | wild-type | repletion | 13.48 | 15.73 | 1.12 |
| *racH*<sup>-</sup> | wild-type | repletion | 16.07 | 7.14 | 0 |
| *atg1*<sup>-</sup> | wild-type | repletion | 53.73 | 14.93 | 4.48 |
| All events considered |  |  |  |  |  |
| *D. discoideum* | *M. marinum* | Condition | Late events (after 10 h.p.i) (%) |  |  |
|  |  |  | Non-lytic release | Lytic release | Bacteria digestion |
| wild-type | wild-type | repletion | 12.75 | 27.45 | 0.98 |
| wild-type | ΔRD1 | repletion | 11.32 | 2.83 | 0.94 |
| *wshA*<sup>-</sup> | wild-type | repletion | 34.83 | 29.21 | 5.62 |
| *racH*<sup>-</sup> | wild-type | repletion | 25 | 51.79 | 0 |
| *atg1*<sup>-</sup> | wild-type | repletion | 2.99 | 23.88 | 0 |
| Reuptakes excluded |  |  |  |  |  |
| *D. discoideum* | *M. marinum* | Condition | Early events (up to 10 h. post starvation or mock) (%) |  |  |
|  |  |  | Non-lytic release | Lytic release | Bacteria digestion |
| wild-type | wild-type | repletion | 31.67 | 11.67 | 3.33 |
| wild-type | wild-type | starvation | 68.18 | 3.41 | 1.14 |
| *wshA*<sup>-</sup> | wild-type | starvation | 4.84 | 11.29 | 0 |
| *racH*<sup>-</sup> | wild-type | starvation | 23.33 | 6.67 | 1.67 |
| *atg1*<sup>-</sup> | wild-type | starvation | 13.51 | 2.70 | 0 |
| Reuptakes excluded |  |  |  |  |  |
| *D. discoideum* | *M. marinum* | Condition | Late events (after 10 h. post starvation or mock) (%) |  |  |
|  |  |  | Non-lytic release | Lytic release | Bacteria digestion |
| wild-type | wild-type | repletion | 15 | 35 | 3.33 |
| wild-type | wild-type | starvation | 13.64 | 13.64 | 0 |
| *wshA*<sup>-</sup> | wild-type | starvation | 56.45 | 24.19 | 3.23 |
| *racH*<sup>-</sup> | wild-type | starvation | 36.67 | 31.67 | 0 |
| *atg1*<sup>-</sup> | wild-type | starvation | 8.11 | 75.68 | 0 |

**Supplementary Table 3.**
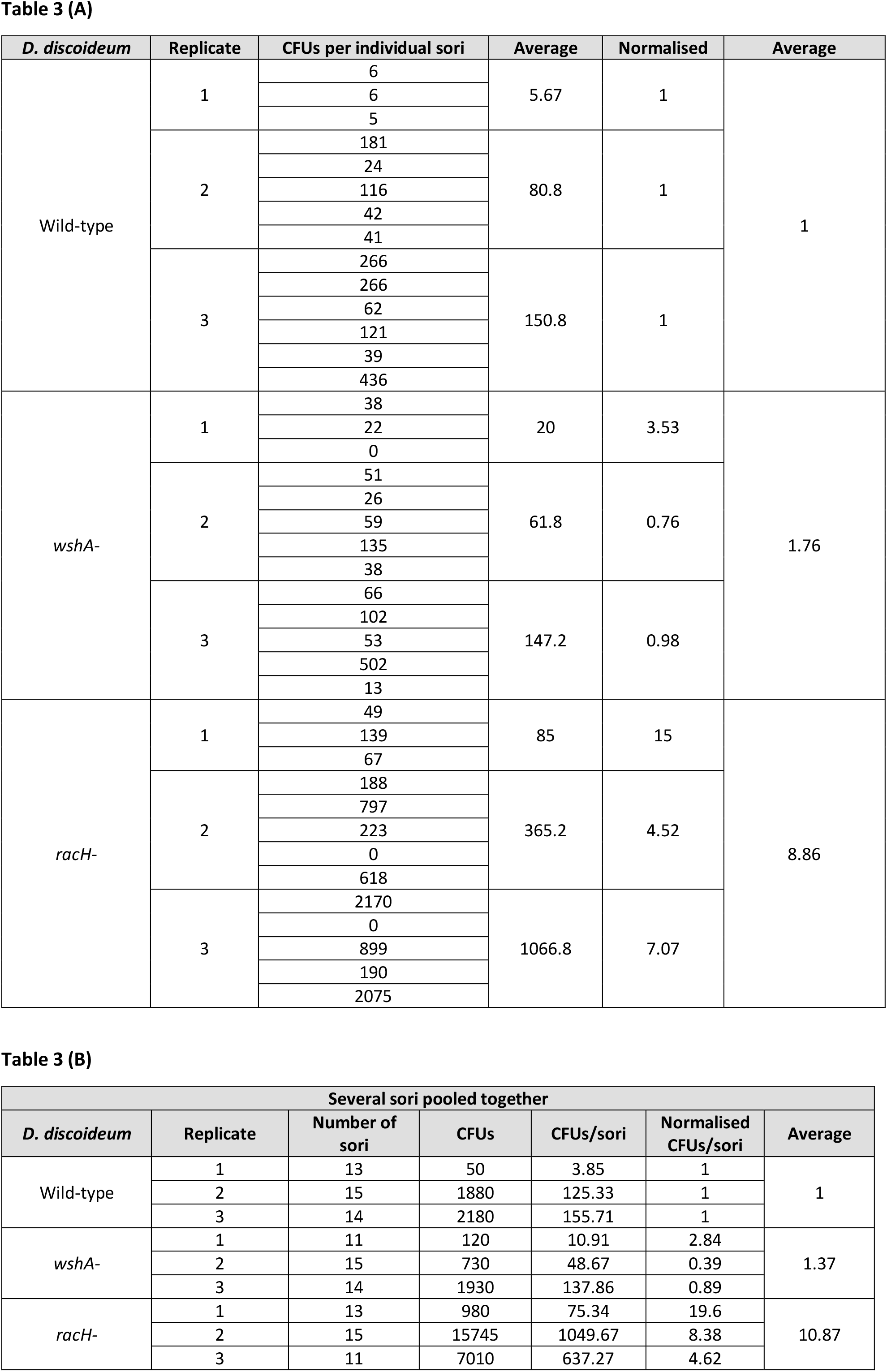
Summary table of the sterility assay results. Number of sori and CFUs quantified during the sterility assay. Data complementary to Fig 7C (Table 3A) and 7D (Table 3B).

**Movie S1. Starvation leads to infection curing.** At 6 hpi, *D. discoideum* infected cells were covered with a thin film of Sorensen buffer and imaged at 25° C every 10 minutes for 24 hours using a fluorescence widefield-microscope (*D. discoideum* visible in the brightfield images, *M. marinum* in green). 4 fields of view were stitched together.

**Movie S2. *D. discoideum* has a good fitness in the InfectChip.** *D. discoideum* cells were trapped in the InfectChip and visualized every 3 min in a spinning disc confocal microscope at 22° C. 6 fields of view were stitched together.

**Movie S3. Long-term single-cell imaging of *D. discoideum* infected with *M. marinum*.** *D. discoideum* cells were infected with mCherry10 expressing *M. marinum* and trapped in the InfectChip for visualization in a spinning disc confocal microscope at 25° C. Brightfield images were taken every 4 min, fluorescent images were taken every 1h.

**Movie S4. Long-term single-cell imaging of *D. discoideum* infected with *M. marinum* ΔRD1.** *D. discoideum* cells were infected with DsRed expressing *M. marinum* ΔRD1 and trapped in the InfectChip for visualization in a spinning disc confocal microscope at 25° C. Brightfield images were taken every 4 min, fluorescent images were taken every 20 min.

**Movie S5. Long-term single-cell imaging of VacA-GFP expressing *D. discoideum* infected with *M. marinum*.** (Left) *D. discoideum* cells expressing GFP-VacA were infected with mCherry10 expressing *M. marinum* and trapped in the InfectChip for visualization in a spinning disc confocal microscope at 25° C. Brightfield images were taken every 4 min, fluorescent images were taken every 1h. (Right) VacA-GFP fluorescence at the MCV was monitored and quantified. Data was normalized to the maximum fluorescence intensity observed.

**Movie S6. Long-term single-cell imaging of *wshA-D. discoideum* infected with *M. marinum*.** *D. discoideum* cells were infected with mCherry10 expressing *M. marinum* and trapped in the InfectChip for visualization in a spinning disc confocal microscope at 25° C. Brightfield images were taken every 4 min, fluorescent images were taken every 1h.

**Movie S7. Long-term single-cell imaging of *racH-D. discoideum* infected with *M. marinum*.** *D. discoideum* cells were infected with mCherry10 expressing *M. marinum* and trapped in the InfectChip for visualization in a spinning disc confocal microscope at 25° C. Brightfield images were taken every 4 min, fluorescent images were taken every 1h.

**Movie S8. Long-term single-cell imaging of *atg1-D. discoideum* infected with *M. marinum*.** *D. discoideum* cells were infected with mCherry10 expressing *M. marinum* and trapped in the InfectChip for visualization in a spinning disc confocal microscope at 25° C. Brightfield images were taken every 4 min, fluorescent images were taken every 1h.

## REFERENCES

Ackermann, M., 2015. A functional perspective on phenotypic heterogeneity in microorganisms. Nat Rev Microbiol 13, 497–508.

Arafah, S., Kicka, S., Trofimov, V., Hagedorn, M., Andreu, N., Wiles, S., Robertson, B., Soldati, T., 2013. Setting up and monitoring an infection of Dictyostelium discoideum with mycobacteria. Methods Mol Biol 983, 403–417.

Baker, J., Brown, R.W., 1994. Trojan Horses of the microbial world: protozoa and the survival of bacterial pathogens in the environment. Microbiology 140, 1253–1259.

Barisch, C., Lopez-Jimenez, A.T., Soldati, T., 2015. Live imaging of Mycobacterium marinum infection in Dictyostelium discoideum. Methods Mol Biol 1285, 369–385.

Benabentos, R., Hirose, S., Sucgang, R., Curk, T., Katoh, M., Ostrowski, E.A., Strassmann, J.E., Queller, D.C., Zupan, B., Shaulsky, G., Kuspa, A., 2009. Polymorphic members of the lag gene family mediate kin discrimination in Dictyostelium. Curr Biol 19, 567–572.

Boulais, J., Trost, M., Landry, C.R., Dieckmann, R., Levy, E.D., Soldati, T., Michnick, S.W., Thibault, P., Desjardins, M., 2010. Molecular characterization of the evolution of phagosomes. Mol Syst Biol 6, 423.

Bozzaro, S., Buracco, S., Peracino, B., 2013. Iron metabolism and resistance to infection by invasive bacteria in the social amoeba Dictyostelium discoideum. Front Cell Infect Microbiol 3, 50.

Bozzaro, S., Eichinger, L., 2011. The Professional Phagocyte Dictyostelium discoideum as a Model Host for Bacterial Pathogens. Current Drug Targets 12, 942–954.

Brock, D.A., Read, S., Bozhchenko, A., Queller, D.C., Strassmann, J.E., 2013. Social amoeba farmers carry defensive symbionts to protect and privatize their crops. Nat Commun 4, 2385.

Cardenal-Munoz, E., Arafah, S., Lopez-Jimenez, A.T., Kicka, S., Falaise, A., Bach, F., Schaad, O., King, J.S., Hagedorn, M., Soldati, T., 2017. Mycobacterium marinum antagonistically induces an autophagic response while repressing the autophagic flux in a TORC1- and ESX-1-dependent manner. PLoS Pathog 13, e1006344.

Carnell, M., Zech, T., Calaminus, S.D., Ura, S., Hagedorn, M., Johnston, S.A., May, R.C., Soldati, T., Machesky, L.M., Insall, R.H., 2011. Actin polymerization driven by WASH causes V-ATPase retrieval and vesicle neutralization before exocytosis. J Cell Biol 193, 831–839.

Carroll, P., Schreuder, L.J., Muwanguzi-Karugaba, J., Wiles, S., Robertson, B.D., Ripoll, J., Ward, T.H., Bancroft, G.J., Schaible, U.E., Parish, T., 2010. Sensitive detection of gene expression in mycobacteria under replicating and non-replicating conditions using optimized far-red reporters. PLoS One 5, e9823.

Chen, G., Zhuchenko, O., Kuspa, A., 2007. Immune-like phagocyte activity in the social amoeba. Science 317, 678–681.

Cirillo, J.D., Falkow, S., Bermudez, L.E., Tompkins, L.S., 1997. Interaction of Mycobacterium avium with Environmental Amoebae Enhances Virulence. Infection and Immunity 65, 3759–3767.

Cosma, C.L., Humbert, O., Ramakrishnan, L., 2004. Superinfecting mycobacteria home to established tuberculous granulomas. Nat Immunol 5, 828–835.

Cosma, C.L., Humbert, O., Sherman, D.R., Ramakrishnan, L., 2008. Trafficking of superinfecting Mycobacterium organisms into established granulomas occurs in mammals and is independent of the Erp and ESX-1 mycobacterial virulence loci. J Infect Dis 198, 1851–1855.

Cosson, P., Lima, W.C., 2014. Intracellular killing of bacteria: is Dictyostelium a model macrophage or an alien? Cell Microbiol 16, 816–823.

Cosson, P., Soldati, T., 2008. Eat, kill or die: when amoeba meets bacteria. Curr Opin Microbiol 11, 271–276.

Delince, M.J., Bureau, J.B., Lopez-Jimenez, A.T., Cosson, P., Soldati, T., McKinney, J.D., 2016. A microfluidic cell-trapping device for single-cell tracking of host-microbe interactions. Lab Chip 16, 3276–3285.

Dhar, N., McKinney, J., Manina, G., 2016. Phenotypic Heterogeneity in Mycobacterium tuberculosis. Microbiol Spectr 4.

Dinh, C., Farinholt, T., Hirose, S., Zhuchenko, O., Kuspa, A., 2018. Lectins modulate the microbiota of social amoebae. Science 361, 402–406.

DiSalvo, S., Haselkorn, T.S., Bashir, U., Jimenez, D., Brock, D.A., Queller, D.C., Strassmann, J.E., 2015. Burkholderia bacteria infectiously induce the protofarming symbiosis of Dictyostelium amoebae and food bacteria. Proc Natl Acad Sci U S A 112, E5029–5037.

Escoll, P., Rolando, M., Gomez-Valero, L., Buchrieser, C., 2013. From amoeba to macrophages: exploring the molecular mechanisms of Legionella pneumophila infection in both hosts. Curr Top Microbiol Immunol 376, 1–34.

Freeze, H., Loomis, W.F., 1977. Isolation and characterization of a component of the surface sheath of Dictyostelium discoideum. J Biol Chem 252, 820–824.

Gao, L.Y., Guo, S., McLaughlin, B., Morisaki, H., Engel, J.N., Brown, E.J., 2004. A mycobacterial virulence gene cluster extending RD1 is required for cytolysis, bacterial spreading and ESAT-6 secretion. Mol Microbiol 53, 1677–1693.

Gerisch, G., 1968. Cell aggregation and differentiation in Dictyostelium. Curr Top Dev Biol 3, 157–197.

Gerstenmaier, L., Pilla, R., Herrmann, L., Herrmann, H., Prado, M., Villafano, G.J., Kolonko, M., Reimer, R., Soldati, T., King, J.S., Hagedorn, M., 2015. The autophagic machinery ensures nonlytic transmission of mycobacteria. Proc Natl Acad Sci U S A 112, E687–692.

Green, D.R., Oguin, T.H., Martinez, J., 2016. The clearance of dying cells: table for two. Cell Death Differ 23, 915–926.

Hagedorn, M., Rohde, K.H., Russell, D.G., Soldati, T., 2009. Infection by tubercular mycobacteria is spread by nonlytic ejection from their amoeba hosts. Science 323, 1729–1733.

Hagedorn, M., Soldati, T., 2007. Flotillin and RacH modulate the intracellular immunity of Dictyostelium to Mycobacterium marinum infection. Cell Microbiol 9, 2716–2733.

Harms, A., Maisonneuve, E., Gerdes, K., 2016. Mechanisms of bacterial persistence during stress and antibiotic exposure. Science 354.

Hirose, S., Benabentos, R., Ho, I., Kuspa, A., Shaulsky, G., 2011. Self-recognition in social amoebae is mediated by allelic pairs of tiger genes. Science 333, 467–470.

Hosseini, R., Lamers, G.E., Soltani, H.M., Meijer, A.H., Spaink, H.P., Schaaf, M.J., 2016. Efferocytosis and extrusion of leukocytes determine the progression of early mycobacterial pathogenesis. J Cell Sci 129, 3385–3395.

Huang, J., Brumell, J.H., 2009. Autophagy in immunity against intracellular bacteria. Current Topics in Microbiology and Immunology 335, 189–215.

Journet, A., Chapel, A., Jehan, S., Adessi, C., Freeze, H., Klein, G., Garin, J., 1999. Characterization of *Dictyostelium discoideum* cathepsin D. Molecular cloning, gene disruption, endo-lysosomal localization and sugar modifications. J. Cell Sci. 112, 3833–3843.

Kang, C.M., Nyayapathy, S., Lee, J.Y., Suh, J.W., Husson, R.N., 2008. Wag31, a homologue of the cell division protein DivIVA, regulates growth, morphology and polar cell wall synthesis in mycobacteria. Microbiology 154, 725–735.

King, J., Gueho, A., Hagedorn, M., Gopaldass, N., Leuba, F., Soldati, T., Insall, R., 2013. wAsH is required for lysosomal recycling and efficient autophagic and phagocytic digestion. Mol Biol Cell. 24, 2714–2726.

Kolonko, M., Geffken, A.C., Blumer, T., Hagens, K., Schaible, U.E., Hagedorn, M., 2014. WASH-driven actin polymerization is required for efficient mycobacterial phagosome maturation arrest. Cell Microbiol 16, 232–246.

Konijn, T.M., Van de Meene, J.G.C., Bonner, J.T., Barkley, D.S., 1967. The acrasin activity of adenosine-3’,5’-cyclic phosphate. Proc Natl Acad Sci USA. 58, 1152–1154.

Lamrabet, O., Mba Medie, F., Drancourt, M., 2012a. Acanthamoeba polyphaga-enhanced growth of Mycobacterium smegmatis. PLoS One 7, e29833.

Lamrabet, O., Merhej, V., Pontarotti, P., Raoult, D., Drancourt, M., 2012b. The genealogic tree of mycobacteria reveals a long-standing sympatric life into free-living protozoa. PLoS One 7, e34754.

Lardy, B., Bof, M., Aubry, L., Paclet, M.H., Morel, F., Satre, M., Klein, G., 2005. NADPH oxidase homologs are required for normal cell differentiation and morphogenesis in Dictyostelium discoideum. Biochim Biophys Acta 1744, 199–212.

Lee, E., Knecht, D.A., 2002. Visualization of actin dynamics during macropinocytosis and exocytosis. Traffic 3, 186–192.

Lelong, E., Marchetti, A., Gueho, A., Lima, W.C., Sattler, N., Molmeret, M., Hagedorn, M., Soldati, T., Cosson, P., 2011. Role of magnesium and a phagosomal P-type ATPase in intracellular bacterial killing. Cell Microbiol 13, 246–258.

Loomis, W.F., 2008. cAMP oscillations during aggregation of Dictyostelium. Adv Exp Med Biol 641, 39–48.

Loomis, W.F., 2014. Cell signaling during development of Dictyostelium. Dev Biol 391, 1–16.

Loomis, W.F., 2015. Genetic control of morphogenesis in Dictyostelium. Dev Biol 402, 146–161.

Lopez-Jimenez, A.T., Cardenal-Munoz, E., Leuba, F., Gerstenmaier, L., Barisch, C., Hagedorn, M., King, J.S., Soldati, T., 2018. The ESCRT and autophagy machineries cooperate to repair ESX-1-dependent damage at the Mycobacterium-containing vacuole but have opposite impact on containing the infection. PLoS Pathog 14, e1007501.

Mahamed, D., Boulle, M., Ganga, Y., Mc Arthur, C., Skroch, S., Oom, L., Catinas, O., Pillay, K., Naicker, M., Rampersad, S., Mathonsi, C., Hunter, J., Wong, E.B., Suleman, M., Sreejit, G., Pym, A.S., Lustig, G., Sigal, A., 2017. Intracellular growth of Mycobacterium tuberculosis after macrophage cell death leads to serial killing of host cells. Elife 6.

Marchetti, A., Lelong, E., Cosson, P., 2009. A measure of endosomal pH by flow cytometry in Dictyostelium. BMC Res Notes 2, 7.

Matz, C., Kjelleberg, S., 2005. Off the hook--how bacteria survive protozoan grazing. Trends Microbiol 13, 302–307.

Mba Medie, F., Ben Salah, I., Henrissat, B., Raoult, D., Drancourt, M., 2011. Mycobacterium tuberculosis complex mycobacteria as amoeba-resistant organisms. PLoS One 6, e20499.

Molmeret, M., Horn, M., Wagner, M., Santic, M., Abu Kwaik, Y., 2005. Amoebae as training grounds for intracellular bacterial pathogens. Appl Environ Microbiol 71, 20–28.

Ochiail, H., Stadler, J., Westphal, M., Wagle, G., Merkl, R., Gerisch, G., 1982. Monoclonal antibodies against contact sites A of Dictyostelium discoideum: detection of modifications of the glycoprotein in tunicamycintreated cells. The EMBOJounal 1, 1011–1016,.

Poff, K.L., Butler, W.L., W.F., L., 1973. Light-Induced Absorbance Changes Associated with Phototaxis in Dictyostelium. Proc. Nat. Acad. Sci. USA 70, 813–816.

Raper, K.B., 1935. Dictyostelium discoideum, a new species of slime mold from decaying forest leaves. Journal of Agricultural Research 50, 135–147.

Salah, I.B., Ghigo, E., Drancourt, M., 2009. Free-living amoebae, a training field for macrophage resistance of mycobacteria. Clin Microbiol Infect 15, 894–905.

Santi, I., Dhar, N., Bousbaine, D., Wakamoto, Y., McKinney, J.D., 2013. Single-cell dynamics of the chromosome replication and cell division cycles in mycobacteria. Nat Commun 4, 2470.

Shu, L., Brock, D.A., Geist, K.S., Miller, J.W., Queller, D.C., Strassmann, J.E., DiSalvo, S., 2018. Symbiont location, host fitness, and possible coadaptation in a symbiosis between social amoebae and bacteria. Elife 7.

Simeone, R., Bobard, A., Lippmann, J., Bitter, W., Majlessi, L., Brosch, R., Enninga, J., 2012. Phagosomal rupture by Mycobacterium tuberculosis results in toxicity and host cell death. PLoS Pathog 8, e1002507.

Siu, C.H., Sriskanthadevan, S., Wang, J., Hou, L., Chen, G., Xu, X., Thomson, A., Yang, C., 2011. Regulation of spatiotemporal expression of cell-cell adhesion molecules during development of Dictyostelium discoideum. Dev Growth Differ 53, 518–527.

Smith, E.W., Lima, W.C., Charette, S.J., Cosson, P., 2010. Effect of starvation on the endocytic pathway in Dictyostelium cells. Eukaryot Cell 9, 387–392.

Solomon, J.M., Leung, G.S., Isberg, R.R., 2003. Intracellular Replication of Mycobacterium marinum within Dictyostelium discoideum: Efficient Replication in the Absence of Host Coronin. Infection and Immunity 71, 3578–3586.

Somesh, B.P., Neffgen, C., Iijima, M., Devreotes, P., Rivero, F., 2006. Dictyostelium RacH regulates endocytic vesicular trafficking and is required for localization of vacuolin. Traffic 7, 1194–1212.

Stinear, T.P., Seemann, T., Harrison, P.F., Jenkin, G.A., Davies, J.K., Johnson, P.D., Abdellah, Z., Arrowsmith, C., Chillingworth, T., Churcher, C., Clarke, K., Cronin, A., Davis, P., Goodhead, I., Holroyd, N., Jagels, K., Lord, A., Moule, S., Mungall, K., Norbertczak, H., Quail, M.A., Rabbinowitsch, E., Walker, D., White, B., Whitehead, S., Small, P.L., Brosch, R., Ramakrishnan, L., Fischbach, M.A., Parkhill, J., Cole, S.T., 2008. Insights from the complete genome sequence of Mycobacterium marinum on the evolution of Mycobacterium tuberculosis. Genome Res 18, 729–741.

Strassmann, J.E., 2016. Kin Discrimination in Dictyostelium Social Amoebae. J Eukaryot Microbiol 63, 378–383.

Tosetti, N., Croxatto, A., Greub, G., 2014. Amoebae as a tool to isolate new bacterial species, to discover new virulence factors and to study the host-pathogen interactions. Microb Pathog 77, 125–130.

Van der Henst, C., Scrignari, T., Maclachlan, C., Blokesch, M., 2016. An intracellular replication niche for Vibrio cholerae in the amoeba Acanthamoeba castellanii. ISME J 10, 897–910.

Veltman, D.M., Akar, G., Bosgraaf, L., Van Haastert, P.J., 2009. A new set of small, extrachromosomal expression vectors for Dictyostelium discoideum. Plasmid 61, 110–118.

Wang, M., Aerts, R.J., Spek, W., Schaap, P., 1988. Cell cycle phase in Dictyostelium discoideum is correlated with the expression of cyclic AMP production, detection, and degradation. Involvement of cyclic AMP signaling in cell sorting. Dev Biol 125, 410–416.

Weijer, C.J., 2004. Dictyostelium morphogenesis. Curr Opin Genet Dev 14, 392–398.

Zhang, X., Soldati, T., 2013. Detecting, visualizing and quantitating the generation of reactive oxygen species in an amoeba model system. J Vis Exp, e50717.

Zhang, X., Zhuchenko, O., Kuspa, A., Soldati, T., 2016. Social amoebae trap and kill bacteria by casting DNA nets. Nat Commun 7, 10938.

